# Organ-Specific Long-Read Transcriptome Assembly of *Stenochlaena palustris* and Annotation of Anthocyanin Biosynthesis Genes

**DOI:** 10.1101/2025.07.30.667584

**Authors:** Maria Stefanie Dwiyanti, Della Rahmawati, Maria Dewi Puspitasari Tirtaningtyas Gunawan Puteri, Deshika Panapitiya, Yanetri Asi Nion, Elza Wijaya, Filiana Santoso

## Abstract

**Background:** *Stenochlaena palustris* is valued as a vegetable and medicinal fern native to Southeast Asia; however, it remains largely underrepresented in genomic studies. People in Kalimantan (Indonesia) collect young leaves and fronds from wild populations for use as vegetables or medicines to treat conditions, such as ulcers, stomachaches, fever, diarrhoea, and skin infections. The young leaves and fronds of *S. palustris* contain flavonoids, polyphenols, and anthocyanins. Here, we present a high-quality organ-specific transcriptome assembly of *S. palustris* based on long-read RNA sequencing of young leaves.

**Results:** The de novo assembly yielded 47,759 transcripts, with an N50 of 1,524 bp and a BUSCO completeness of 66.6%, consistent with organ-specific transcriptomes. Functional annotation identified key structural and regulatory genes involved in anthocyanin biosynthesis, including genes for chalcone synthase (CHS) and dihydroflavonol 4-reductase (DFR). We further analysed the expression of the selected *CHS* and *DFR* genes via qRT-PCR of three phenotypically contrasting young leaf samples. Although no strong correlation was observed between gene expression levels and anthocyanin pigmentation, the results suggest that complex regulation involves post-transcriptional control or developmental timing.

**Conclusions:** This study provides the first long-read transcriptomic resource for *S. palustris* and valuable data for future investigations of secondary metabolism and gene regulation in ferns. Our findings complement broader fern transcriptome studies by offering tissue-specific resolution and a focused view of pigment biosynthesis.

## BACKGROUND

*Stenochlaena palustris*, a climbing fern of the Blechnaceae family, is a naturally distributed plant in South Asia, Southeast Asia, Australia, and Polynesia [1,2]. It is known by several local names, such as kelakai/kalakai in Kalimantan (Indonesia); lemidin, midin, or paku midin in Malaysia; and diliman or hagnaya in the Philippines [3,4,5,6]. In these regions, young leaves and fronds of *S. palustris* are collected from wild habitats and consumed as a vegetable [3,4,5,6]. In addition to its culinary uses, *S. palustris* has been traditionally used for its medicinal properties for treating conditions, such as ulcers, stomachaches, fever, diarrhoea, and skin infection [4,5,6]. It also treats anaemia, promotes breast milk production, aids recovery after childbirth, prevents diabetes, and reduces antimicrobial activity [4,5,6]. Additionally, *S. palustris* is a hardy plant capable of growing in challenging environments, such as acidic peatlands. In Kalimantan, it is commonly found in various habitats, including acidic swamp areas, riversides, natural and secondary forests, palm oil plantations, and residential zones [1,6,7]. It is also known as a pioneer plant, which can grow first after land disturbances, such as peatland forest fires, thus creating conditions conducive to the establishment of other plant species [8]. These properties render *S. palustris* a valuable source of vegetables and medicines, particularly for the people of Kalimantan.

Studies have shown that fronds and young leaves of *S. palustris* possess antioxidant properties. Water extracts of fronds contain anthocyanins, polyphenols, and hydroxycinnamic acids, which contribute to their antioxidant activity [9]. Young leaves of *S. palustris* are either red– or green-coloured, indicating the differences in anthocyanin content. Young red leaves turn green when they mature [6]. Two independent studies identified the α-glucosidase inhibitor activity of *S. palustris* [9,10]. This inhibitory activity may be attributed to hydroxycinnamic acids or astragalin [9,10]. Astragalin is kaempferol 3-O-β-glucopyranoside, a glucosylated form of kaempferol. It is found in various plant species, including persimmon leaves, lotus leaves, green tea seeds, and roots of *Astragalus membranaceus* [11].

Understanding the regulation of flavonoid and anthocyanin biosynthesis is crucial for further phytochemical-related research on *S. palustris.* Flavonoid biosynthesis begins with the conversion of phenylalanine to the flavonoid precursor *p*-coumaroyl-CoA through a series of enzymatic steps [12]. The key enzymes involved in this process are phenylalanine ammonia-lyase (PAL), cinnamate 4-hydroxylase (C4H), and 4-coumaroyl-CoA ligase (4CL) [12]. *p*-Coumaroyl-CoA is converted to naringenin chalcone by chalcone synthase (CHS). This intermediate is further converted into naringenin by chalcone isomerase (CHI). The addition of a hydroxyl group at the 3-position of naringenin results in the formation of dihydrokaempferol, a process catalysed by flavanone 3-hydroxylase (F3H). Dihydrokaempferol is subsequently converted into kaempferol by flavonol synthase (FLS). Kaempferol is glycosylated by UDP-dependent glycosyltransferases (UGT) to form astragalin. Additionally, dihydrokaempferol serves as a precursor for anthocyanins, with several enzymes involved in the conversion, including dihydroflavonol 4-reductase (DFR), flavonoid 3□-hydroxylase (F3□H), flavonoid 3□,5□-hydroxylase (F3□5□H), anthocyanidin synthase (ANS), flavonoid 3-O-glucosyltransferase (F3GT), and flavonoid 5-O-glucosyltransferase (F5GT).

The growing interest in flavonoid and anthocyanin biosynthesis in ferns has led to the identification of related genes. The first step in flavonoid biosynthesis, which is the conversion of *p*-coumaryl CoA to naringenin chalcone by CHS, is often rate-limiting. CHS genes were isolated from the ferns *Dryopteris fragrans* (GenBank accession numbers KF530802.1, KP420005.1, and KP420004.1), *D. erythrosora* (GenBank accession number KJ135628.1), and *Ceratopteris thalictroides* (GenBank: JX027616.1). As CHS can exist as a single copy or form a multigene family, the actual number of CHS-expressing genes in ferns may be higher [13]. DFR is the key enzyme involved in anthocyanin biosynthesis. DFR genes have been identified and characterised in water ferns (*Azolla filiculoides*) and *D. erythrosora.* Chen et al. [14] isolated two DFR genes from *D. erthyrosora*, namely *DeDFR1* (GenBank: MK920230) and *DeDFR2* (GenBank: MK920231). Based on a GenBank search, Chen et al. [14] deposited three other DFR genes: MK920232, MK920233, and MK920234. In this study, these genes were designated as *DFR3*, *DFR4*, and *DFR5*. Two DFR gene families were identified in *A. filiculoides*: *DFR1* consisting of Azfi_s0035.g025620 and Azfi_s0245.g059984 and *DFR2* with Azfi_s0008.g011655 and Azfi_s0008.g011657. [15,16].

Limited genomic information on *S. palustris* hinders the characterisation of flavonoid biosynthesis regulation. Currently, only three transcriptomic datasets are publicly available in the National Center for Biotechnology Information Sequence Read Archive (NCBI SRA) database (https://www.ncbi.nlm.nih.gov/sra; last accessed March 20^th^, 2025). The three datasets were generated from samples from China and Singapore [17,18,19]. Shen et al. [18] generated the transcriptome data for *S. palustris* sporophyll and trophophyll tissues. Sequencing was performed using the Illumina HiSeq 2500 system. *De novo* assembly produced 58,416 contigs with 945.83 bp average length and 48.15% GC content [18]. Ali et al. [19] determined the transcriptome profile of *S. palustris* from various organs, such as sterile and fertile leaves, petioles, rhizomes, and roots. Sequencing was performed using the Illumina NovaSeq 6000 system. The *de novo* assembly produced 54,843 contigs with 832.1 bp average length and 45.25% GC content [19]. The *de novo* assembly of the transcriptome data produced by Ali et al. [19] is publicly available (https://conekt.plant.tools/, last accessed March 21^st^, 2025).

In this study, we used the Nanopore long-read sequencing platform to obtain transcriptome data from leaf samples collected from Palangka Raya, Indonesia. The data obtained in the present study can support genomic research on *S. palustris* by providing additional information that complements previously published data. In the present study, we identified genes potentially involved in flavonoid and anthocyanin biosynthesis, particularly those encoding CHS and DFR.

## METHODS

### Sample collection, RNA extraction, and sequencing

Young green nonfibrous sterile leaves from *S. palustris* naturally grown in the University of Palangka Raya (UPR), Indonesia (2°12′57.2″S, 113°54′04.3″E) were collected and immediately stored in RNAlater solution (ThermoFisher Scientific, Japan). Plant material from a location in UPR (2°12′48.4″S, 113°54′02.8″E) had been previously identified at the Herbarium Bogoriense, Research Center for Biology, Cibinong, Indonesia (No. 252/IPH.1.01/If.07/II/2019).

The samples were stored at 4 °C until RNA extraction. RNA was extracted from leaves using 2% cetyltrimethylammonium bromide (CTAB) and 4% polyvinylpyrrolidone (PVP) extraction buffer, following the protocol of Kiss et al. [20], with several modifications. Leaf tissue (5 cm) was ground to a fine powder in liquid nitrogen using a mortar and a pestle. A preheated extraction buffer (6 mL) was added to the powder, and the mixture was homogenised to remove clumps. The mixture was transferred to six 2 mL tubes. The tubes were then incubated at 65 °C for 10 min. Seven hundred microlitres of chloroform:isoamyl alcohol (24:1) was added to each tube. The tubes were vortexed and centrifuged at 15,000 rpm for 10 min at 4 °C. The upper phase was transferred to a new 1.5 mL tube, and an equal volume of chloroform:isoamyl alcohol (24:1) was added. The addition of chloroform:isoamyl alcohol was repeated three times. The upper phase was then mixed with 500 μL 8M LiCl. The mixture was incubated at 4 °C overnight. The following day, the solution was centrifuged at 15,000 rpm for 45 min at 4 °C. The RNA pellet was washed with 50 µL ice-cold ethanol (80% v/v) and then centrifuged at 15,000 rpm for 5 min at 4 °C. The supernatant was carefully removed using a pipette, and the tubes were centrifuged and then dried at 37 °C to remove any residual ethanol. The RNA pellets were resuspended in 270 μL RNase-free water. Two microlitres of DNase I (Nippongene, Japan), 27 μL DNase I buffer, and 1 μL recombinant RNase inhibitor (Takara Shuzo, Kyoto, Japan) were added to the RNA solution and then incubated at 37 °C for 30 min. DNase was removed using phenol:chloroform:isoamyl alcohol (25:24:1). RNA was precipitated in 99.5% ethanol overnight at –30 °C then centrifuged at 15,000 rpm for 45 min at 4 °C. The RNA pellet was washed using 100 μL cold 80% EtOH, followed by another centrifugation at 15,000 rpm for 5 min at 4 °C. Residual ethanol was removed using a pipette, and the pellet was air-dried at room temperature. The pellet was eluted in 50 μL RNase/DNase-free water.

RNA quantity was determined using NanoDrop spectrophotometer (Thermo Fisher Scientific, Wilmington, DE, USA) and Qubit (Thermo Fisher Scientific, Japan). RNA quality was assessed via 1% gel electrophoresis, and Bioanalyzer (Agilent Technologies, Santa Clara, CA, USA). Transcriptome sequencing was performed on the PromethION platform using a cDNA-PCR Barcoding Kit 24 V14 (SQK-PCR114.24) at GeneBay Inc., Japan. Base calling was performed using the Dorado base-call server (version 7.4). A high-accuracy model was used to generate sequence reads after filtering. To ensure high-quality data in the downstream analysis, low-quality and short reads were removed, and barcode sequences were trimmed using Porechop 0.2.4 [21], with the remove-middle option applied to eliminate barcodes embedded within the reads.

### *De novo* assembly, reduce redundancy, and assembly quality assessment

*De novo* sequence assembly of reads that passed filtering was performed using RNA-Bloom2 version 2.0.1, with the default parameters [22]. To reduce redundancy, the contigs were clustered based on sequence similarity using CD-HIT-EST in the CD-HIT package version 4.8.1 [23], with the following parameters: similarity threshold (-c 0.90) and word size threshold (-n 8). The resulting contigs were discarded if 1) read coverage <1 based on read mapped to each contig by minimap [24], 2) sequences are predicted from non-fern organisms by the evaluation conducted using NCBI Foreign Contamination Screen (NCBI FCS) provided in Galaxy (https://usegalaxy.org, last accessed March 27^th^, 2025), 3) barcode adapters remain in the sequences by filtering manually before uploading to NCBI TSA database, and 4) GC% of the contig is more than 60% or less than 40%. The assembly quality was evaluated using Benchmarking Universal Single-Copy Orthologs v5 (BUSCO v5) [25,26] to assess the completeness of the assembly. Online BUSCO analysis used Embryophyta from the OrthoDB v10 ortholog sets (https://gvolante.riken.jp [27]). The final dataset was examined for the contig length distribution, including the minimum, maximum, and average values, and the %GC content was assessed using QUAST [28].

### Identification of open reading frame

Open reading frames (ORFs) for each contig were predicted using ORFipy version 0.0.4 [29]. ORFipy was run using the following parameters: start ATG to search for ORFs beginning at the common start codon ATG and Table 1 standard genetic code, which is applicable for most organisms. Option –longest was also added to filter and output the largest ORFs. The largest ORFs predicted for each contig were assembled into a dataset to predict gene function.

**Table 1.**
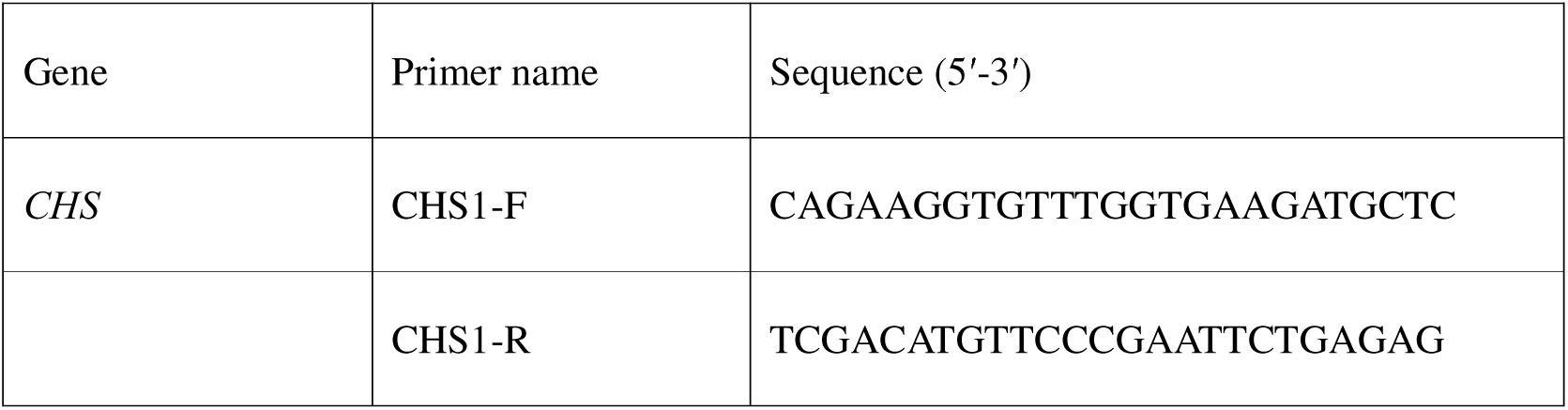
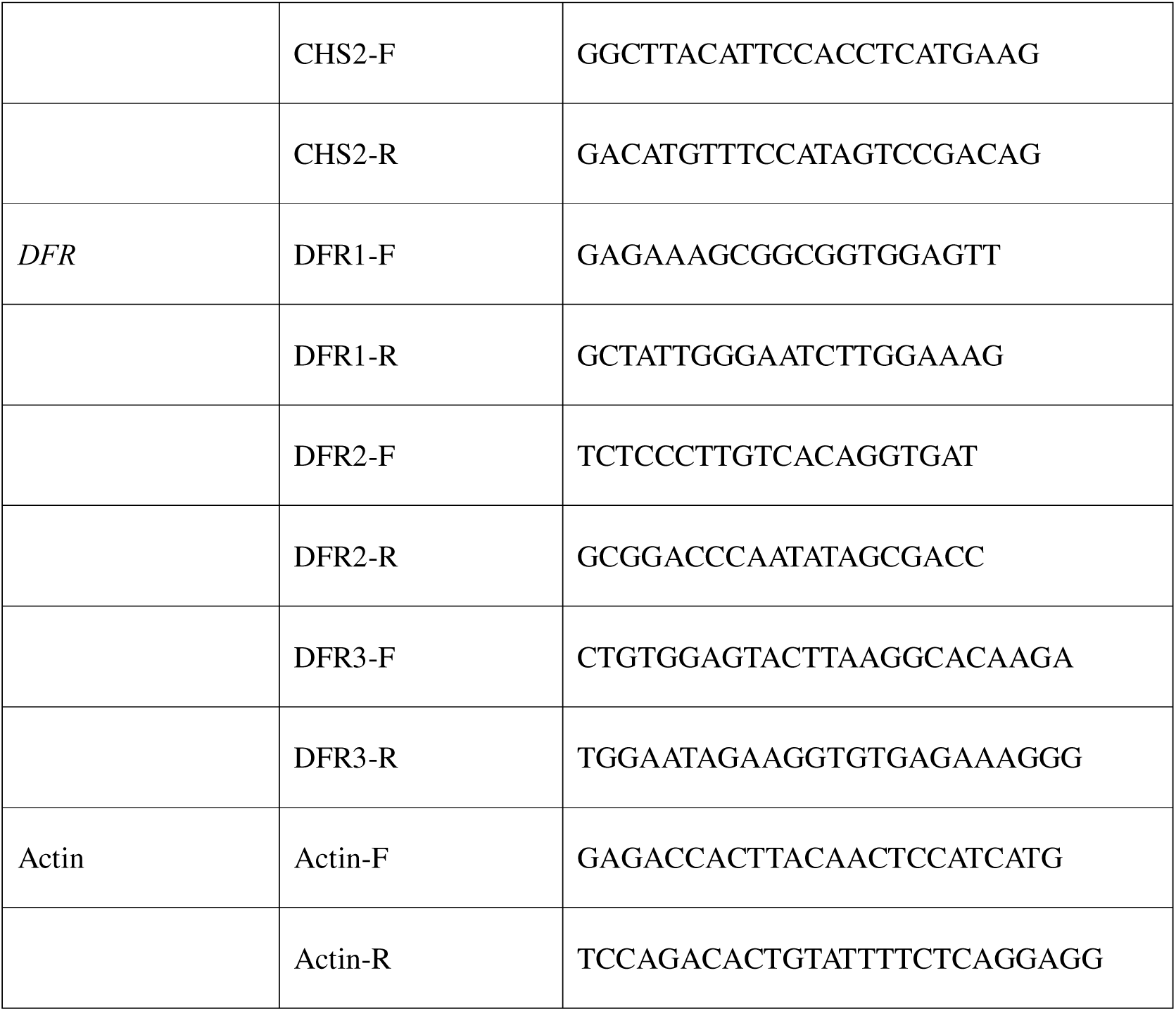
Primers used for quantitative real-time PCR.

### Gene functional annotation using eggNOG-mapper v2

Functional annotation was performed using the eggNOG-mapper v2 web version (http://eggnog-mapper.embl.de [30,31])by uploading the longest contigs obtained from the CD-HIT-EST analysis. The input was nucleotide sequences, whereas the parameters used were genomic data, gene prediction method: BLASTX-like, allow frameshifts (Diamond –frameshift option), database eggnog 5, minimum hit e-value 0.001, minimum hit bit-score 60, percentage identity 40%, minimum% of query coverage 20%, and minimum% of subject coverage 20%. For further analysis, another filtering was performed to retain only the contigs that matched “max_annotation_level” to “Streptophyta” or “Viridiplantae”. KEGG orthology (KO) and Clusters of Orthologous Groups (COG) distributions were used for the final output contigs.

### Mining flavonoid biosynthesis genes

Based on the output from the eggNOG-mapper v2, genes potentially involved in flavonoid biosynthesis were identified and compiled (Figure 3, Additional File 1). Multiple sequence alignment and phylogenetic tree analysis were performed to define gene clustering. For CHS, only 16 genes with lengths of > 800 bp were included in the analysis, whereas all 11 DFR genes were included. In addition to *S. palustris* genes, known CHS and DFR genes from other ferns were included for comparison. The CHS genes included were *D. fragrans* (GenBank: KF530802.1, KP420005.1, KP420004.1), *D. erythrosora* (GenBank: KJ135628.1), *C. thalictroides* (GenBank: JX027616.1), whereas the DFR genes included were *D. erythrosora* (MK920230.1, MK920231.1, MK920232.1, MK920233.1, MK920234.1) and *A. filiculoides* (Azfi_s0035.g025620, Azfi_s0245.g059984, Azfi_s0008.g011655, Azfi_s0008.g011657). Multiple sequence alignment was performed using MAFFT version 7 with the L-INS-I algorithm (https://mafft.cbrc.jp/alignment/server/index.html [32]). A phylogenetic tree was constructed using the neighbour-joining method with the Jukes-Cantor substitution model and 1000-times bootstrap replicates provided on the MAFFT web server. The phylogenetic tree was edited using Archaeopteryx.js, which is provided on the same server.

### Gene expression analysis via quantitative real-time PCR

To confirm the expression of genes involved in flavonoid biosynthesis, quantitative real-time PCR (qRT-PCR) was conducted using RNA extracted from young leaves collected from natural populations at three locations in Central Kalimantan: the UPR, Kalampangan, and Tjilik Riwut (Figure 5A). Sampling locations were the UPR (2°12′57.2″S, 113°54′04.3″E), Tjilik Riwut (2°14′02.8″S, 113°57′09.1″E), and Kalampangan (2°16′44.8″S, 114°00′30.3″E). The leaves from the UPR site were green, whereas those from Kalampangan and Tjilik Riwut were red. The supernatant phase of the water-methanol-chloroform extract of the UPR leaves was clear, whereas that of Kalampangan and Tjilik Riwut leaves was pink, probably due to presence of anthocyanins [33]. The samples were kept fresh in an icebox and were lyophilised for 24 hours before storage at –80 °C. RNA extraction and DNase I treatment were performed as previously described. RNA quantity and quality were measured using the NanoDrop spectrophotometer.

cDNA synthesis was performed as follows: a mixture consisting of 200 ng total RNA, 1 µL 50 μM oligo (dT)_20_ primer (Toyobo, Osaka, Japan), 1 mol dNTP mixture (Takara Shuzo, Kyoto, Japan) was denatured at 65 °C for 5 min and then immediately put on ice. 5× RT buffer (4 μL), RNase inhibitor (1 μL), and M-MLV RT (1 μL, Invitrogen, Tokyo, Japan) were added to the mixture. The cDNA synthesis reaction was then carried out at 42 °C for 60 min followed by a denaturation at 70 °C for 15 minutes and a cooling down step at 10 °C, infinite time.

A 10 µL reaction mixture was prepared for qRT-PCR: 5 μL SSoAdvanced Universal SYBR Green Supermix (Bio-Rad, Hercules, CA, USA), 2 μL cDNA, and 250 nM forward and reverse primers. The qRT-PCR conditions were as follows: initial denaturation at 95 °C for 30 s, followed by 40 cycles each at 95 °C for 30 s, 58 °C for 15 s, and 72 °C for 20 s. Melting curve analysis ranged from 56 to 95 °C. To examine the specificity of the qRT-PCR amplification, temperature was increased by 0.5 °C every 10 s after completion of the amplification cycles. The gene expression level of each sample was calculated and compared with the expression level of the internal control gene actin. Primers designed for the selected genes are listed in Table 1. For gene expression analysis, primers were designed targeting the conserved regions. qRT-PCR was performed using the Bio-Rad CFX Connect machine at the Faculty of Agriculture, Hokkaido University. Three biological replicates were prepared.

## RESULTS

### Long-read nanopore sequencing and *de novo* assembly results

Long-read nanopore sequencing generated 15,632,749 raw reads with a total nucleotide length of 13,033,432,831 bp. After filtering, 14,913,665 reads, with an average length of 682.5 bp, were retained for *de novo* assembly. The *de novo* assembly produced 112,160 contigs. The reduced-redundancy step using CD-HIT-EST organised the 112,160 contigs into 74,467 clusters, from which the longest contig in each cluster was selected as the representative and used for further analysis. After further filtering based on read coverage, absence of adapter sequences, and GC%, a final set of 47,759 contigs was obtained. The minimum length was 500 bp, the maximum length was 11,260 bp, and the average length was 1328.8 bp (Figure 1C). The GC content was 45.15%. Completeness scores based on BUSCO assessment were 66.6% complete (44.9% single-copy and 21.7% duplicated), 6.2% fragmented, and 27.2% missing (Figure 1C).

**Figure 1.**
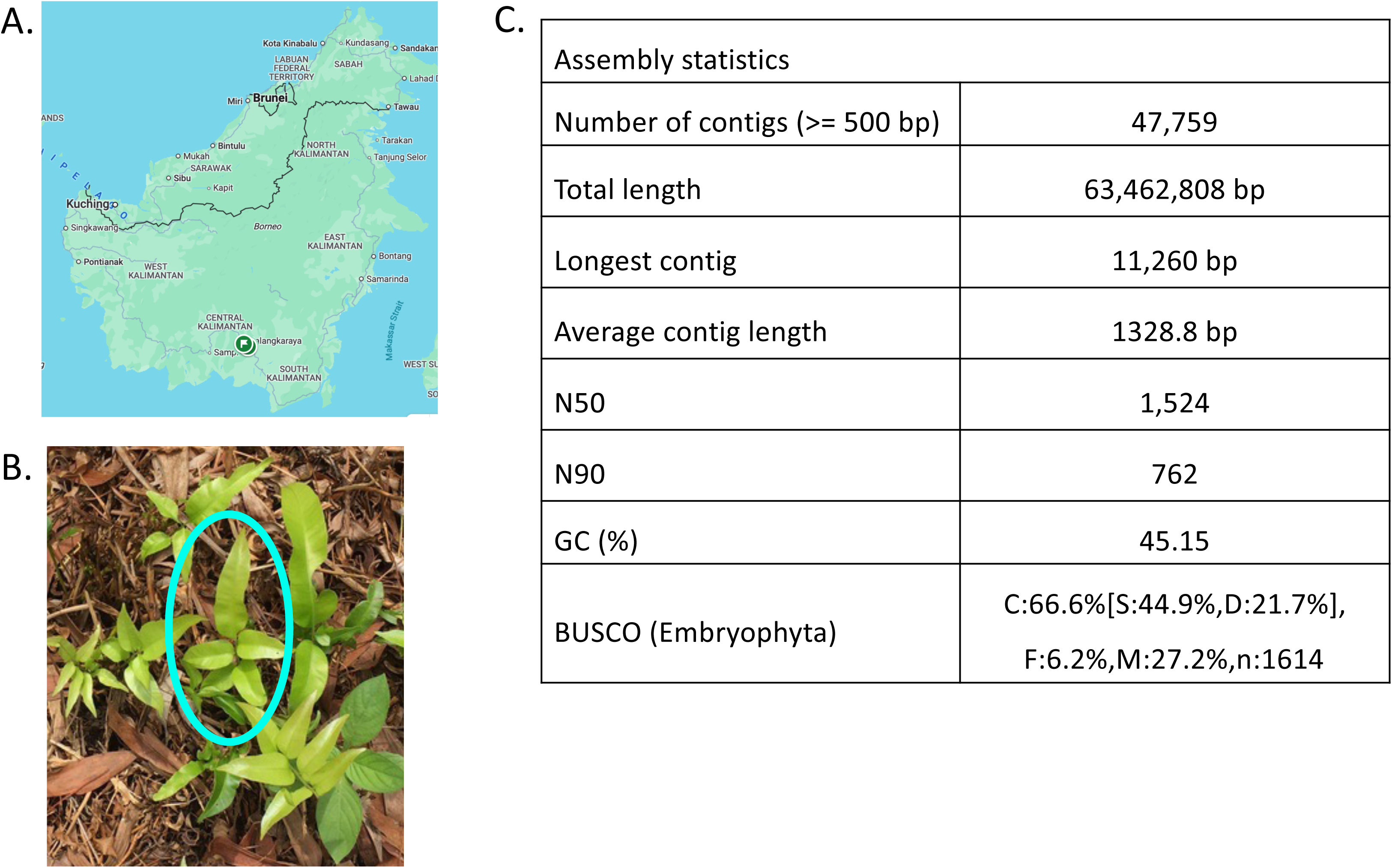
A) Sampling location on Kalimantan Island. The map was sourced from Google Maps (https://maps.google.com/. Accessed April 23, 2025). B) Leaf samples used for Nanopore long-read sequencing. C) Assembly statistics.

### Gene annotation based on eggNOG-mapper

The longest ORFs from 47,759 contigs were predicted using ORFipy. The resulting dataset was functionally annotated using eggNOG-mapper. A total of 30,010 contigs were successfully annotated, with the highest annotation level assigned to either *Viridiplantae* or *Streptophyta*. KO terms were assigned to each contig; some contigs had a single KO term, whereas others had multiple terms. Based on the KO classifications, 3,745 contigs were associated with genetic information processing, 1,206 with environmental information processing, 82 with organismal systems, and 5,960 with metabolism (Figure 2A). Among those classified under “Metabolism”, 301 contigs were linked to “biosynthesis of other secondary metabolites”, including genes involved in the biosynthesis of phenylpropanoid, flavone and flavonol, isoflavonoid, flavonoid, and anthocyanin.

**Figure 2.**
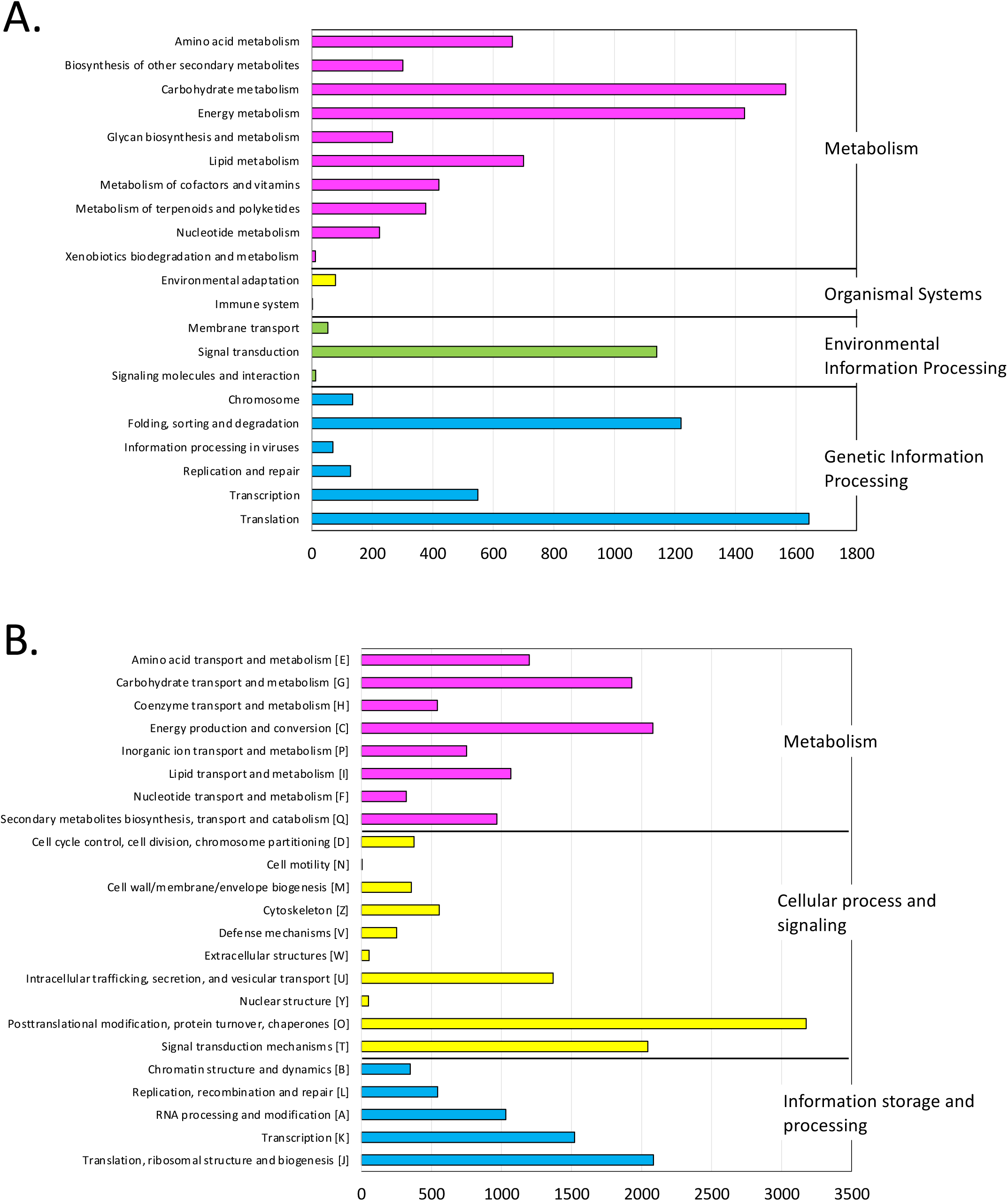
EggNOG-mapper analysis result. Genes were categorised based on A. KEGG orthology groups and B. Clusters of Orthologous Groups. The numbers show the count of genes belonging to each group.

**Figure 3.**
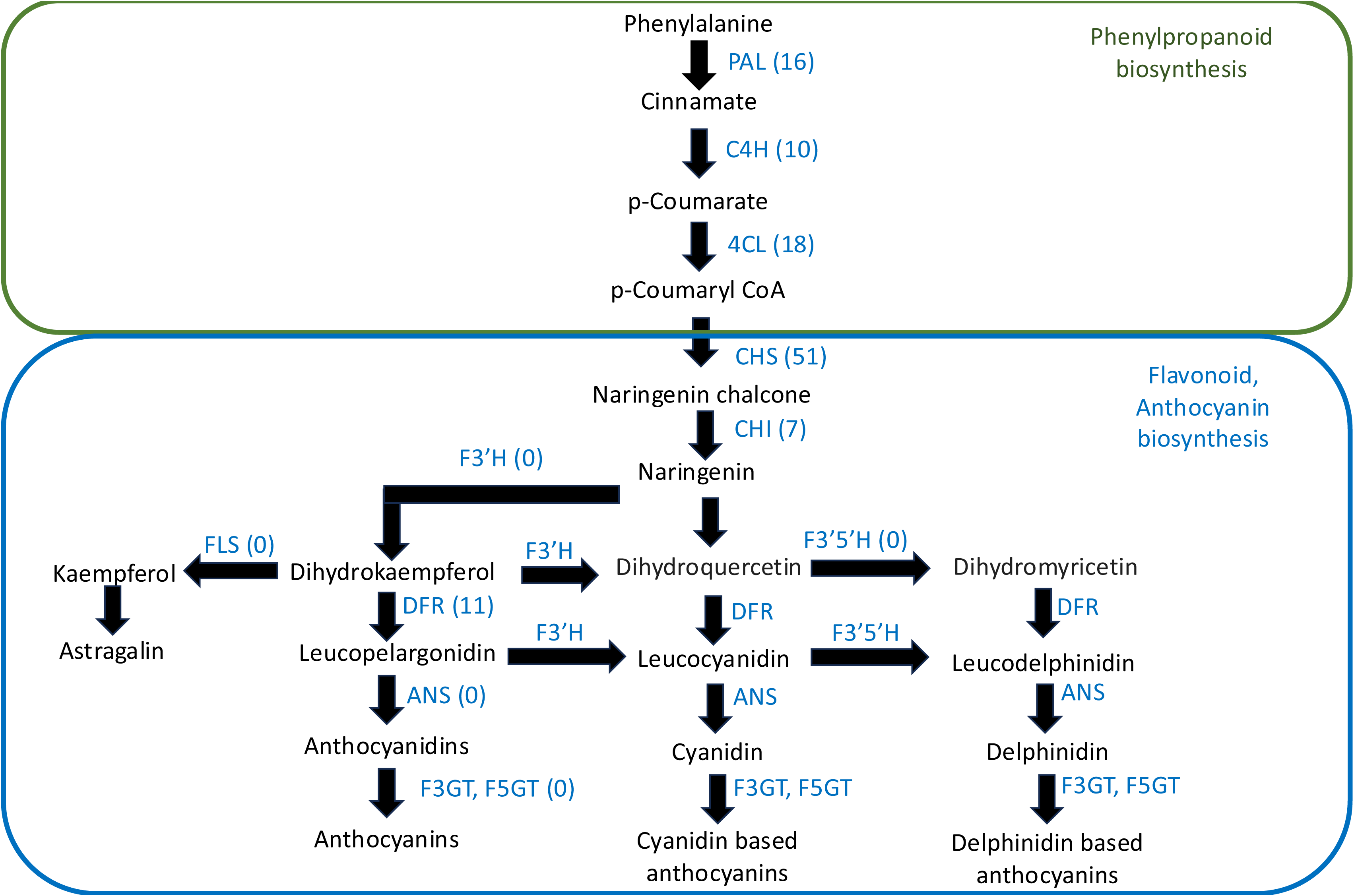
Phenylpropanoid, flavonoid, and anthocyanin pathway for *Stenochlaena palustris*. PAL, phenylalanine ammonia-lyase; C4H, cinnamate 4-hydroxylase; 4CL, 4-coumarate:CoA ligase; CHS, chalcone synthase; CHI, chalcone isomerase; F3H, flavanone 3-hydroxylase; DFR, dihydroflavonol 4-reductase; ANS, anthocyanidin synthase; F3GT, flavonoid 3-*O*-glucosyltransferase; F5GT, flavonoid 5-*O*-glucosyltransferase; FLS, flavonol synthase; F3□H, flavonoid 3□-hydroxylase; F3□5□H, flavonoid 3□,5□-hydroxylase; UGT, UDP-dependent glycosyltransferase. The numbers in brackets show the number of genes identified for each enzyme.

Based on COG annotations, 5,531 contigs were assigned to information storage and processing, 8,238 to cellular processes and signaling, and 8,858 to metabolism (Figure 2B). Within the metabolism category, the contigs related to energy production and conversion (C), carbohydrate transport and metabolism (G), and amino acid transport and metabolism (E) were the most abundant. Additionally, 967 contigs were classified as associated with secondary metabolite biosynthesis, transport, and catabolism (Q) (Figure 2B).

### Mining flavonoid biosynthesis-related genes

Genes or contigs encoding enzymes for phenylpropanoid and flavonoid biosynthesis were identified based on eggNOG-mapper annotation. There were 16 genes for PAL, 10 for C4H, 18 for 4CL, 51 for CHS, 7 for CHI, and 11 for DFR (Figure 3). No genes were identified for F3′H, F3′5′H, FLS, ANS, F3GT, and F5GT.

The selected *CHS* genes identified in this study were aligned with *CHS* genes from the ferns *D. fragrans*, *D. erythrosora*, and *C. thalictroides*. Phylogenetic analysis using the neighbour-joining method clustered *CHS* genes into three major groups, designated CHS1, CHS2, and CHS3 (Figure 4; Additional File 2-4). The CHS1 cluster included three genes: rb_8683, rb_49489, and rb_54131 (Additional File 2). Compared with rb_49489, rb_54131 had a 264-bp deletion and a 1-bp insertion that shifted the predicted stop codon downstream. rb_8683 showed a 99-bp deletion but retained the original stop codon position (Additional File 2). CHS2 comprised rb_4659, rb_49361, and rb_103035 (Additional File 3). rb_103035 lacked the first 306 bp whereas rb_4659 and rb_49361 differed by four single nucleotide polymorphisms. The CHS3 group included four genes: rb_7618, rb_74283, rb_15005, and rb_52876 (Additional File 4). While their 5′ regions are highly conserved, variations in nucleotide composition and deletions in the 3′ regions resulted in differences in amino acid sequences and stop codon positions (Additional File 4).

**Figure 4.**
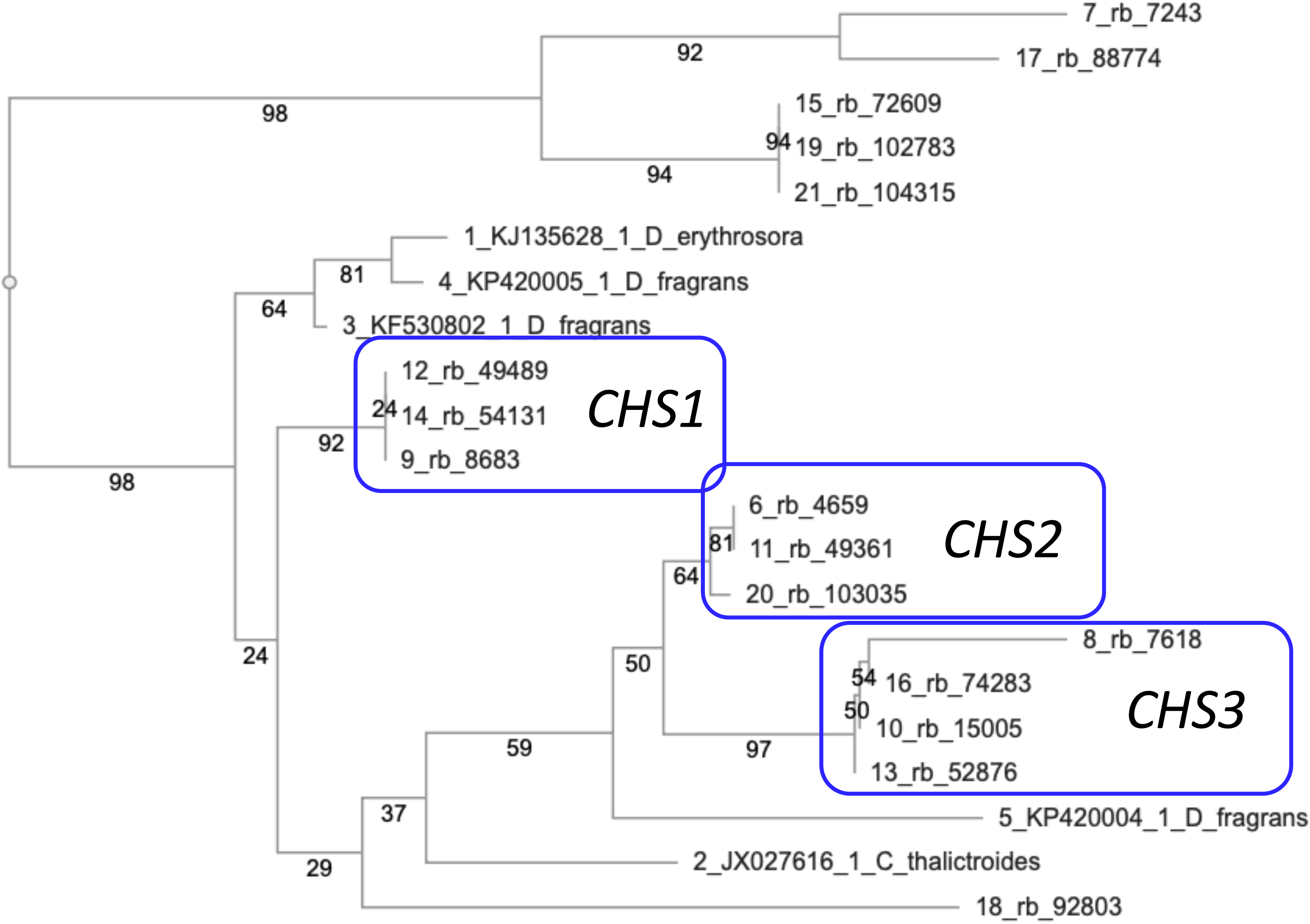
Neighbor-joining tree of chalcone synthase (*CHS*) genes. Fifty-one CHS-coding genes obtained in this study and *CHS* genes from *Dryopteris fragrans* (KF530802.1), *D. erythrosora* (KJ135628.1) and *Ceratopteris thalictroides* (JX027616.1) are included. The tree was built using 1,000 bootstrap replicates.

**Figure 5.**
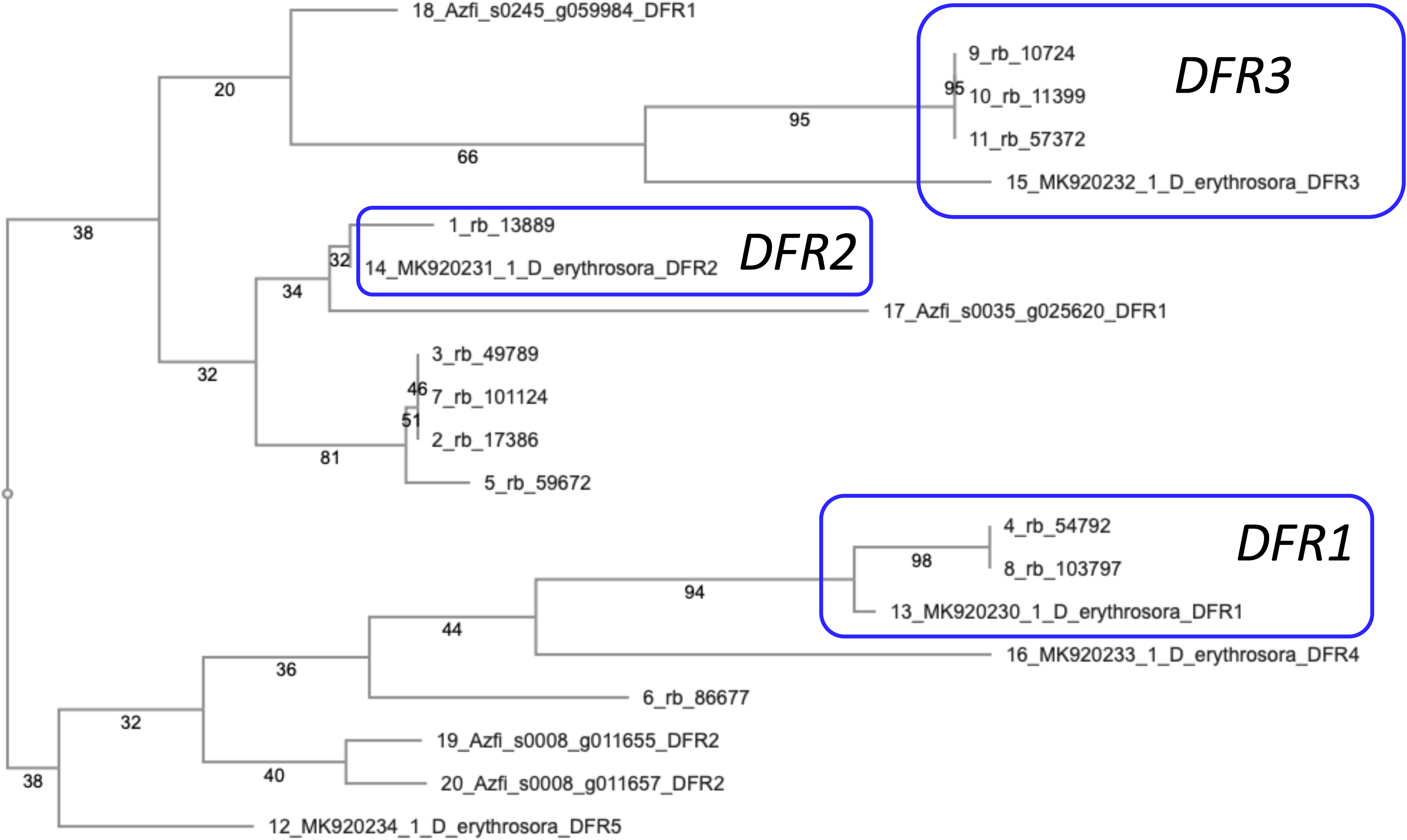
Neighbor-joining tree of dihydroflavonol 4-reductase (*DFR*) genes. DFR-coding genes obtained in this study and five *DFR* genes from *Dryopteris erythrosora,* and four *DFR* genes from *Azolla filiculoides* are included. The tree was built using 1,000 bootstrap replicates.

To investigate the relationships among *DFR* genes, 11 putative *DFR* sequences, along with *DFR* genes from *D. erythrosora* and *A. filiculoides*, were analysed using the neighbour-joining method (Figure 5). rb_54792 and rb_103797 clustered with *D. erythrosora DFR1* and were classified as part of the DFR1 group. Compared with rb_103797, rb_54792 contained a 117-bp deletion at the 5′ end and an additional 146-bp deletion that resulted in an upstream shift of the stop codon (Additional File 5). rb_13889 grouped with *D. erythrosora DFR2* and was designated as part of the DFR2 group (Additional File 6). Three genes, rb_10724, rb_11399, and *rb_57372*, clustered with *D. erythrosora DFR3* and were assigned to the DFR3 group. Within this group, rb_10724 exhibited a 315-bp deletion at the 5′ end compared with rb_11399, whereas rb_57372 had a 90-bp insertion-deletion (indel) at the same region. Additionally, rb_57372 showed a 114-bp indel and several nucleotide variations in the 3′ region, leading to an earlier stop codon (Additional File 7). The remaining four genes formed a distinct clade, separate from the known *DFR* genes of *A. filiculoides* and *D. erythrosora*, suggesting the presence of novel *DFR* variants.

To asses gene expression via qRT-PCR, gene-specific primers were designed based on the conserved regions of *CHS1*, *CHS2*, *DFR1*, *DFR2*, and *DFR3*. Expression of these genes was confirmed in young leaf tissues collected from populations in the UPR, Kalampangan, and Tjilik Riwut (Figure 6A). qRT-PCR further validated the expression of both *CHS* and *DFR* across all populations (Figure 6B). However, the variations in leaf and leaf extract colouration observed among the UPR, Kalampangan, and Tjilik Riwut genotypes did not correspond to differences in *CHS* or *DFR* gene expression levels.

**Figure 6.**
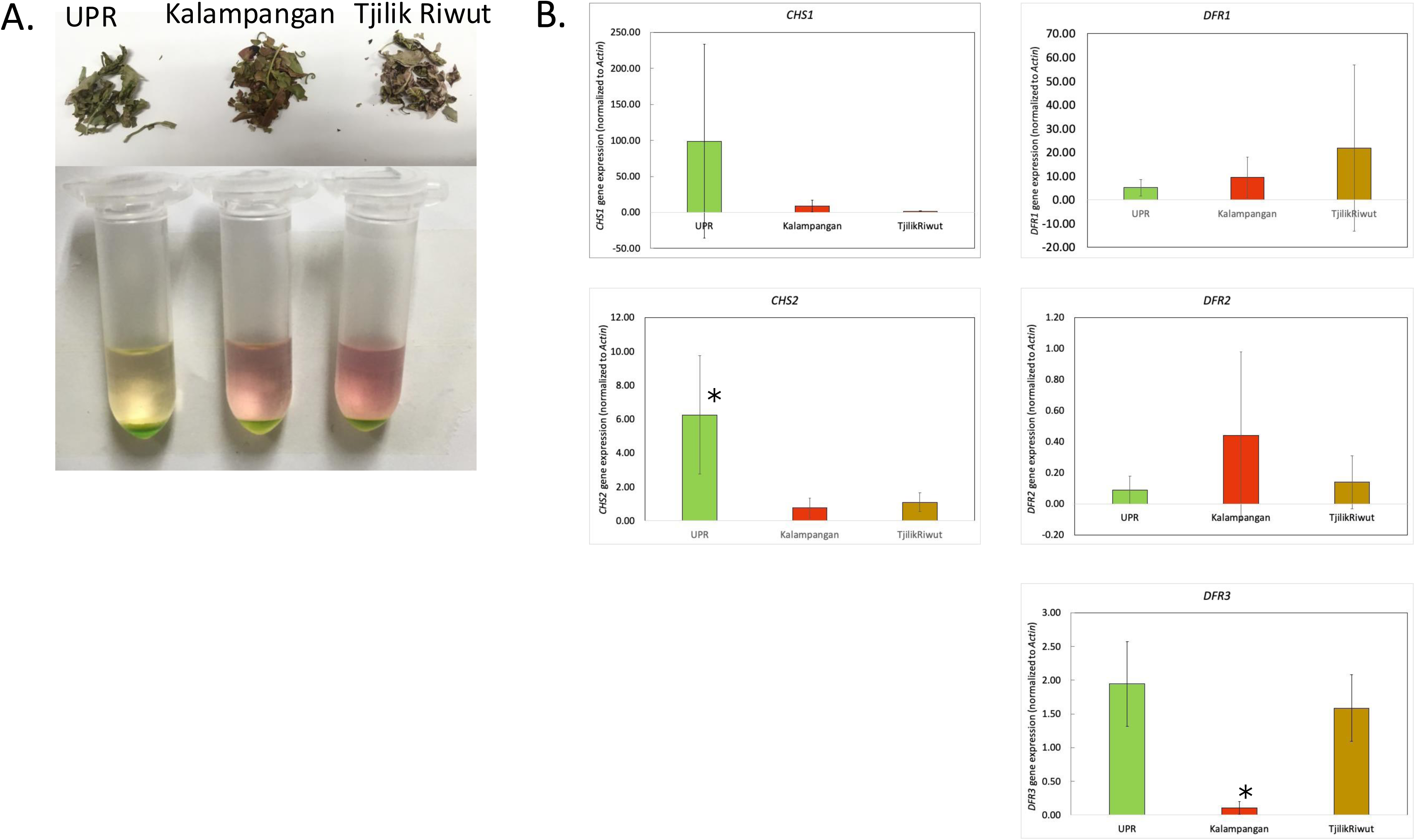
A. Young leaf samples from the University of Palangka Raya (UPR), Tjilik Riwut, and Kalampangan sites, and water-methanol-chloroform extract from the corresponding samples. Sampling locations are the UPR (2°12′57.2″S, 113°54′04.3″E), Tjilik Riwut (2°14′02.8″S, 113°57′09.1″E), and Kalampangan (2°16′44.8″S, 114°00′30.3″E). The map was sourced from Google Maps, 2025. B. Expression levels of *CHS1, CHS2, DFR1*, *DFR2*, and *DFR3* genes in young leaves of the UPR, Kalampangan, and Tjilik Riwut samples. Expression levels were normalised to actin. Three replicates were prepared for each sample. Asterisks indicate a statistically significant difference at *p*<0.05.

## DISCUSSION

The transcriptome profile of *S. palustris* was generated via Nanopore long-read sequencing. *De novo* assembly yielded 47,759 contigs (Figure 1). However, the completeness of the assembly, as evaluated by BUSCO using the Embryophyta dataset, was 66.6% (Figure 1). This relatively low score may be attributed to the use of RNA extracted solely from young leaf tissues, which likely did not capture the full transcriptome complexity of *S. palustris* [26].

Putative genes encoding enzymes involved in flavonoid and anthocyanin biosynthesis were successfully identified. The long-read transcriptome enabled the differentiation of highly similar isoforms of *CHS* and *DFR*, highlighting the advantage of this technology for distinguishing closely related sequences in *de novo* assemblies. The expression of *CHS* and *DFR* genes was confirmed in young leaves through qRT-PCR. However, because the primers targeted conserved regions, the expression of individual gene isoforms could not be determined, warranting further investigation. Interestingly, the gene expression levels of *CHS* and *DFR* did not correlate with visible leaf pigmentation, suggesting that post-transcriptional mechanisms or metabolic regulation influenced pigment accumulation (Figure 6B).

Samples were collected from natural populations at three sites with distinct soil types: UPR, Kalampangan, and Tjilik Riwut. The observed variation in leaf colouration may be due to underlying genetic differences or environmental factors [16,34]. Previous studies have shown that environmental conditions influence the accumulation of flavonoids and phenolic compounds in fern leaves. Particularly, salinity and full sunlight did not alter total polyphenol content in *D. erythrosora* or *A. nipponicum* var. “Red Beauty” but increased the total flavonoid content in *D. erythrosora* [16,34]. Similarly, high light intensity and low temperature enhanced anthocyanin accumulation in *A. filiculoides* fronds [16]. Future studies should assess the expression of flavonoid and anthocyanin biosynthesis genes across various tissues, developmental stages, and controlled environmental conditions to better understand the regulatory mechanisms underlying flavonoid and anthocyanin production in *S. palustris*.

Notably, genes encoding key enzymes in the flavonoid biosynthetic pathway, such as *FLS*, *ANS*, *F3GT*, and *F5GT*, were undetected. This finding aligns with that of Ali et al. [19], who reported that several fern species, including *S. palustris*, lack these genes. Similarly, *FLS* and *ANS* were absent in *C. richardii* [35], although the upstream and downstream genes were present. Although *FLS*-like and *ANS*-like genes have been predicted in *A. filiculoides* [15], BLAST searches of the current transcriptome data did not yield any matches (data not shown). These results suggest that ferns utilise alternative biosynthetic routes or enzymes to fulfil the roles typically performed by the canonical *FLS* and *ANS*.

This transcriptome profiling represents an important step in the characterisation of *S. palustris*. Future studies should focus on obtaining full genome sequences to confirm gene copy numbers, identify genetic markers, and support comparative genomics in ferns. Ultimately, integrated transcriptomic and metabolomic analyses will be essential to fully elucidate the flavonoid and anthocyanin biosynthetic pathways in *S. palustris*.

## CONCLUSIONS

In the present study, we utilised long-read sequencing and *de novo* assembly of the transcriptome profiles of *S. palustris* leaves. The genes involved in flavonoid biosynthesis were also identified. The expression of genes encoding CHS and DFR was confirmed in young leaves of *S. palustris*. These data will facilitate advancements in the genetic and molecular studies of *S. palustris*.

## DECLARATIONS

### Ethics approval and consent to participate

Not applicable.

### Consent for publication

Not applicable.

### Availability of data and materials

The datasets generated and/or analysed during the current study are available in the NCBI Short Read Archive repository under the BioProject ID PRJNA1237869, BioSample accession number SAMN47441734, and SRA accession number SRR32761149 [https://www.ncbi.nlm.nih.gov/bioproject/PRJNA1237869/, https://www.ncbi.nlm.nih.gov/biosample/SAMN47441734/, https://www.ncbi.nlm.nih.gov/sra/SRR32761149]. Contigs resulting from the de novo assembly were deposited in the NCBI Transcriptome Shotgun Assembly Sequence Database under the accession number GLFY00000000.

### Competing interests

The authors declare that they have no competing interests.

### Funding

This study was supported by the Heiwa Nakajima Foundation for MSD.

### Authors’ contributions

MSD conceived the study, performed experiments, and wrote the manuscript. DR collected the samples and edited the manuscript. DP performed the real-time PCR analysis. EW, FS, MDPTGP, and YAN selected the sampling locations, helped with sample collection and preparation before RNA extraction, and edited the manuscript.

## Acknowledgements

We would like to thank Editage (www.editage.jp) for English language editing.

## ADDITIONAL FILES

Additional File 1. List of genes (contigs) annotated as flavonoid biosynthesis genes.

Additional File 2. Alignment of genes in the *CHS1* group. Only coding sequences were used for alignment. Primer binding sites are indicated by blue arrows.

Additional File 3. Alignment of genes in the *CHS2* group. Only coding sequences were used for alignment. Primer binding sites are indicated by blue arrows.

Additional File 4. Alignment of genes in the *CHS3* group. Only coding sequences were used for alignment.

Additional File 5. Alignment of genes in the *DFR1* group. Only coding sequences were used for alignment. Primer binding sites are indicated by blue arrows.

Additional File 6. Alignment of genes in the *DFR2* group. Only coding sequences were used for alignment. Primer binding sites are indicated by blue arrows.

Additional File 7. Alignment of genes in the *DFR3* group. Only coding sequences were used for alignment. Primer binding sites are indicated by blue arrows.

## Notes

### Competing Interest Statement

The authors have declared no competing interest.

