## Additional File 2 for "Organ-Specific Long-Read Transcriptome Assembly of *Stenochlaena palustris* and Annotation of Anthocyanin Biosynthesis Genes"

```
CLUSTAL format alignment by MAFFT (v7.511)


rb_8683     ATGGAAGCTGCTGCTGCTCCTCCTGCCTCTTACAAGCCCTGCCCGCGCAAGATGGAGCGC
rb_49489    ATGGAAGCTGCTGCTGCTCCTCCTGCCTCTTACAAGCCCTGCCCGCGCAAGATGGAGCGC
rb_54131    ATGGAAGCTGCTGCTGCTCCTCCTGCCTCTTACAAGCCCTGCCCGCGCAAGATGGAGCGC
            ************************************************************

rb_8683     GCTGATGGCCCCGCTACGGTGCTCGCTATTGGAACTGCCAACCCGCCTAATGTCTTCCTG
rb_49489    GCTGATGGCCCCGCTACGGTGCTCGCTATTGGAACTGCCAACCCGCCTAATGTCTTCCTG
rb_54131    GCTGATGGCCCCGCTACGGTGCTCGCTATTGGAACTGCCAACCCGCCTAATGTCTTCCTG
            ************************************************************

rb_8683     CAGAGTCAATACCCCGACTTCTACTTCAACATCACCAACAGCAACCACATGGCCGACCTC
rb_49489    CAGAGTCAATACCCCGACTTCTACTTCAACATCACCAACAGCAACCACATGGCCGACCTC
rb_54131    CAGAGTCAATACCCCGACTTCTACTTCAACATCACCAACA--------------------
            ****************************************

rb_8683     AAGGAGAAGTTCCAGCGAATGTGTGACAAGTCAGGGATCACCAAGAGGTACATGTATCTG
rb_49489    AAGGAGAAGTTCCAGCGAATGTGTGACAAGTCAGGGATCACCAAGAGGTACATGTATCTG
rb_54131    ------------------------------------------------------------


rb_8683     AATGAGGAGATACTGAAAGCCAACCCAAGCATGTGCGCATACTGGGAGAAGTCGCTGGAT
rb_49489    AATGAGGAGATACTGAAAGCCAACCCAAGCATGTGCGCATACTGGGAGAAGTCGCTGGAT
rb_54131    ------------------------------------------------------------


rb_8683     GTGAGGCAGGATATGGTGGTGGTGGAGGTGCCCAAGCTGGGCAAGGAGGCAGCTGCCAAG
rb_49489    GTGAGGCAGGATATGGTGGTGGTGGAGGTGCCCAAGCTGGGCAAGGAGGCAGCTGCCAAG
rb_54131    ------------------------------------------------------------


rb_8683     GCCATCAAAGAGTGGGGGCAGCCCAAGTCCAAGATAACCCATCTCATTTTCTGCACCACC
rb_49489    GCCATCAAAGAGTGGGGGCAGCCCAAGTCCAAGATAACCCATCTCATTTTCTGCACCACC
rb_54131    ------------------------------------------------------------


rb_8683     AGTGGCGTTGACATGCCTGGGGCTGATTGGGCACTCACCAAACTTCTTGGGCTCCGACCA
rb_49489    AGTGGCGTTGACATGCCTGGGGCTGATTGGGCACTCACCAAACTTCTTGGGCTCCGACCA
rb_54131    ----GCGTTGACATGCCTGGGGCTGATTGGGCACTCACCAAACTTCTTGGGCTCCGACCA
                ********************************************************

rb_8683     AGTGTGAAGCGACTGATGATGTACCAGCAAGGCTGCTTTGCCGGAGGAACAGTCATGAGA
rb_49489    AGTGTGAAGCGACTGATGATGTACCAGCAAGGCTGCTTTGCCGGAGGAACAGTCATGAGA
rb_54131    AGTGTGAAGCGACTGATGATGTACCAGCAAGGCTGCTTTGCCGGAGGAACAGTCATGAGA
            ************************************************************

rb_8683     ATCGCCAAGGATTTAGCAGAGAACAACAAAGGAGCAAGAGTCCTGGTGGTTTGCAGTGAA
rb_49489    ATCGCCAAGGATTTAGCAGAGAACAACAAAGGAGCAAGAGTCCTGGTGGTTTGCAGTGAA
rb_54131    ATCGCCAAGGATTTAGCAGAGAACAACAAAGGAGCAAGAGTCCTGGTGGTTTGCAGTGAA
            ************************************************************

rb_8683     TTAACTGCGGTTACTTTCCGGGGCCCAAGTGAGACACATCTTGATAGCTTAGTTGGCCAA
rb_49489    TTAACTGCGGTTACTTTCCGGGGCCCAAGTGAGACACATCTTGATAGCTTAGTTGGCCAA
rb_54131    TTAACTGCGGTTACTTTCCGGGGCCCAAGTGAGACACATCTTGATAGCTTAGTTGGCCAA
            ************************************************************

rb_8683     GCATTGTTTGGAGATGGTGCATCTGCAGTGATTATTGGGTCGGATCCTATTCCTCAAGTA
rb_49489    GCATTGTTTGGAGATGGTGCATCTGCAGTGATTATTGGGTCGGATCCTATTCCTCAAGTA
rb_54131    GCATTGTTTGGAGATGGTGCATCTGCAGTGATTATTGGGTCGGATCCTATTCCTCAAGTA
            ************************************************************

rb_8683     GAGAGGCCATGGTTCGAAGTGCACTATGTGGCATCAAACATCTTA---------------
rb_49489    GAGAGGCCATGGTTCGAAGTGCACTATGTGGCATCAAACATCTTACCTGATAGTGACGGA
rb_54131    GAGAGGCCATGGTTCGAAGTGCACTATGTGGCATCAAACATCTTACCTGATAGTGACGGA
            *********************************************

rb_8683     ------------------------------------------------------------
rb_49489    GCCATTGATGGCCACCTGCGTGAGGTCGGCCTAACATTCCATCTGATGAAGGATGTCCCT
rb_54131    GCCATTGATGGCCACCTGCGTGAGGTCGGCCTAACATTCCATCTGATGAAGGATGTCCCT


rb_8683     ------------------------TCTGTTTTGAAAGATTCCTTTCAGAAGGTGTTTGGT
rb_49489    GGCATCATCTCGAAGAACATTGGCTCTGTGTTGAAAGATTCCTTTCAGAAGGTGTTTGGT
rb_54131    GGCATCATCTCGAAGAACATTGGCTCTGTGTTGAAAGATTCCTTTCAGAAGGTGTTTGGT
                                    ***** ******************************

rb_8683     GAAGATGCTCCCTCTTTCAATGACTTGTTCTGGATTGCGCACCCAGGCGGGCCTGCAATT
rb_49489    GAAGATGCTCCCTCTTTCAATGACTTGTTCTGGATTGCGCACCCAGGCGGGCCTGCAATT
rb_54131    GAAGATGCTCCCTCTTTCAATGACTTGTTCTGGATTGCGCACCCAGGCGGGCCTGCAATT
            ************************************************************

rb_8683     CTCGATCAAGTGGAGCAAAAGCTGCAGCTAAAGCCCGAGAAAATGGCGCCAAGTAGGCAT
rb_49489    CTCGATCAAGTGGAGCAAAAGCTGCAGCTAAAGCCCGAGAAAATGGCGCCAAGTAGGCAT
rb_54131    CTCGATCAAGTGGAGCAAAAGCTGCAGCTAAAGCCCGAGAAAATGGCGCCAAGTAGGCAT
            ************************************************************

rb_8683     GTGCTCTCAGAATTCGGGAACATGTCGAGTGCATGTGTAATTTTCATCATGGATCATTTG
rb_49489    GTGCTCTCAGAATTCGGGAACATGTCGAGTGCATGTGTAATTTTCATCATGGATCATTTG
rb_54131    GTGCTCTCAGAATTCGGGAACATGTCGAGTGCATGTGTAATTTTCATCATGGATCATTTG
            ************************************************************

rb_8683     CGCAAGAAATCAGTAGAGCAGAATGCAACAACGACTGGGGAAGGGTACGAGTGGGGGTTG
rb_49489    CGCAAGAAATCAGTAGAGCAGAATGCAACAACGACTGGGGAAGGGTACGAGTGGGGGTTG
rb_54131    CGCAAGAAATCAGTAGAGCAGAATGCAACAACGACTGGGGAAGGGTACGAGTGGGGGTTG
            ************************************************************

rb_8683     CTTCTCGGGTTTGGCCCAGGCCTAACATGT-GAGACGGTGGTTTTGCGAAGTGTTCCGCT
rb_49489    CTTCTCGGGTTTGGCCCAGGCCTAACATGT-GAGACGGTGGTTTTGCGAAGTGTTCCGCT
rb_54131    CTTCTCGGGTTTGGCCCAGGCCTAACATGTGGAGACGGTGGTTTTGCGAAGTGTTCCGCT
            ****************************** *****************************

rb_8683     TGTAGCAAATTAA-----------------------------------------
rb_49489    TGTAGCAAATTAA-----------------------------------------
rb_54131    TGTAGCAAATTAAATAATATGGCAATCAAAAGCAGGTATTCGCAGATGGCATAA
            *************
```
