## Additional File 3 for "Organ-Specific Long-Read Transcriptome Assembly of *Stenochlaena palustris* and Annotation of Anthocyanin Biosynthesis Genes"

CLUSTAL format alignment by MAFFT (v7.511)

```
rb_4659    ATGCCGGTCCCTACTGGTGCTACCTTTCCCCCCTGCGCTCGCAAGATGGAGCGTGCTGAG
rb_49361   ATGCCGGTCCCTACTGGTGCTACCTTTCCCCCCTGCGCTCGCAAGATGGAGCGTGCTGAG
rb_103035  ------------------------------------------------------------


rb_4659    GGGCCTGCTACGGTGCTGGCCATCGGAACTGCGAATCCGCCCAATGCGGTCGACCAGAGT
rb_49361   GGGCCTGCTACGGTGCTGGCCATCGGAACTGCGAATCCGCCCAATGCGGTCGACCAGAGT
rb_103035  ------------------------------------------------------------


rb_4659    ACTTACCCTGACTTCTACTTCAACATCACCAACAGCAATCATATGACTGATCTGAAAGAG
rb_49361   ACTTACCCTGACTTCTACTTCAACATCACCAACAGCAATCATATGACTGATCTGAAAGAG
rb_103035  ------------------------------------------------------------


rb_4659    AAATTCCAGCGCATGTGTGAGAAGTCTGGGATTAAGAAGAGGTACATGTATTTGAATGAG
rb_49361   AAATTCCAGCGCATGTGTGAGAAGTCTGGGATTAAGAAGAGGTACATGTATTTGAATGAG
rb_103035  ------------------------------------------------------------


rb_4659    GAAATCTTGAAGGCCAACCCTAGCATGTGTGCATACTGGGAGAAATCCTTAGATGTGAGA
rb_49361   GAAATCTTGAAGGCCAACCCTAGCATGTGTGCATACTGGGAGAAATCCTTAGATGTGAGA
rb_103035  ------------------------------------------------------------


rb_4659    CAAGACATGGTGGTTGTGGAGGTGCCTAAGCTGGGAAAGGAAGCTGCAGCCAAGGCTATC
rb_49361   CAAGACATGGTGGTTGTGGAGGTGCCTAAGCTGGGAAAGGAAGCTGCAGCCAAGGCTATC
rb_103035  ------ATGGTGGTTGTGGAGGTGCCTAAGCTGGGAAAGGAAGCTGCAGCCAAGGCTATC
                 ******************************************************

rb_4659    AAGGAGTGGGGGCAGCCCAAGTCGAAGATCACTCACCTTATCTTCTGCACCACTAGTGGG
rb_49361   AAGGAGTGGGGGCAGCCCAAGTCGAAGATCACTCACCTTATCTTCTGCACCACTAGTGGG
rb_103035  AAGGAGTGGGGGCAGCCCAAGTCGAAGATCACTCACCTTATCTTCTGCACCACTAGTGGG
           ************************************************************

rb_4659    GTGGATATGCCGGGTGCCGATTGGGCTCTCACGAAGCTGCTGGGGCTGCGCCCCAGTGTC
rb_49361   GTGGATATGCCGGGTGCCGATTGGGCTCTCACGAAGCTGCTGGGGCTGCGCCCCAGTGTC
rb_103035  GTGGATATGCCGGGTGCCGATTGGGCTCTCACAAAGCTGCTGGGGCTGCGCCCCAGTGTC
           ********************************.***************************

rb_4659    AAGAGGCTCATGATGTATCAACAAGGTTGCTTTGCAGGTGGCACAGTTATGAGAATTGCA
rb_49361   AAGAGGCTCATGATGTATCAACAAGGTTGCTTTGCAGGTGGCACAGTTATGAGAATTGCA
rb_103035  AAGAGGCTCATGATGTATCAACAAGGTTGCTTTGCAGGTGGCACAGTTATGAGAATTGCA
           ************************************************************

rb_4659    AAGGATCTGGCGGAAAACAACAAGGGAGCCAGAGTTTTGGTTGTGTGTAGTGAGCTAACT
rb_49361   AAGGATCTGGCGGAAAACAACAAGGGAGCCAGAGTTTTGGTTGTGTGTAGTGAGCTAACT
rb_103035  AAGGATCTGGCGGAAAACAACAAGGGAGCCAGAGTTTTGGTTGTGTGTAGTGAGCTAACT
           ************************************************************

rb_4659    GCTGTGACCTTCCGCGGGCCCAGTGACACACACTTAGACAGTCTTGTTGGGCAAGCTTTG
rb_49361   GCTGTGACCTTCCGCGGGCCCAGTGACACACACTTAGACAGTCTTGTTGGGCAAGCTTTG
rb_103035  GCTGTGACCTTCCGCGGCCCCAGTGACACACACTTAGACAGTCTTGTTGGGCAAGCTTTG
           ***************** ******************************************

rb_4659    TTCGGGGATGGTGCTTCAGCCATCATCGTAGGGGCTGACCCCATCCCAGAAGTAGAGAGG
rb_49361   TTCGGGGATGGTGCTTCAGCCATCATCGTAGGGGCTGACCCCATCCCAGAAGTAGAGAGG
rb_103035  TTCGGGGATGGTGCTTCAGCCATCATCGTAGGGGCTGACCCCATCCCAGAAGTAGAGAGG
           ************************************************************

rb_4659    CCCTGGTTTGAGATTCACTATGTGGCATCCAACATCCTGCCAGATAGCGATGGTGCCATT
rb_49361   CCCTGGTTTGAGATTCACTATGTGGCATCCAACATCCTGCCAGATAGCGATGGTGCCATT
rb_103035  CCCTGGTTTGAGATTCACTATGTGGCATCCAACATCCTGCCGGATAGCGATGGTGCCATT
           *****************************************.******************

rb_4659    GATGGACATCTCCGTGAAGTTGGGCTTACATTCCACCTCATGAAGGATGTGCCAGGGATC
rb_49361   GATGGACATCTCCGCGAAGTTGGGCTTACATTCCACCTCATGAAGGATGTGCCAGGGATC
rb_103035  GATGGACATCTCCGCGAAGTTGGGCTTACATTCCACCTCATGAAGGATGTGCCAGGGATC
           **************.*********************************************

rb_4659    ATTTCTAAGAACATTGGCACTGTATTGAAGGATGCATTTGAGAAGGTCTTTGGCAATGAG
rb_49361   ATTTCTAAGAACATTGGCACTGTATTGAAGGATGCCTTTGAGAAGGTCTTTGGCAATGAG
rb_103035  ATTTCTAAGAACATTGGCACTGTATTGAAGGATGCCTTTGAGAAGGTCTTTGGTAATGAG
           *********************************** *****************.******

rb_4659    GAAGGGGAAGTCCCTAGCTACAATGATGTGTTTTGGATTGCACACCCTGGTGGTCCTGCT
rb_49361   GAAGGGGAAGTCCCTAGCTACAATGATGTGTTTTGGATTGCACACCCTGGTGGTCCTGCC
rb_103035  GAAGGGGAAGTCCCTAGCTACAATGATGTGTTTTGGATTGCACACCCTGGTGGTCCTGCC
           ***********************************************************.

rb_4659    ATTCTAGATCAAGTGGAGCAGAAGTTGCAGCTTAAGCCAGAGAAAATGGCTCCGAGCCGA
rb_49361   ATTCTAGATCAAGTGGAGCAGAAGTTGCAGCTGAAGCCAGAGAAAATGGCTCCGAGCCGA
rb_103035  ATTCTAGATCAAGTGGAGCAGAAGTTGCAGCTGAAGCCAGAGAAAATGGCTCCGAGCCGA
           ******************************** ***************************

rb_4659    CAAGTTCTGTCGGACTATGGAAACATGTCAAGTGCTTGTGTGATTTTCATAATGGACCAC
rb_49361   CAAGTTCTGTCGGACTATGGAAACATGTCAAGTGCTTGTGTGATTTTCATAATGGACCAT
rb_103035  CAAGTTCTGTCGGACTATGGAAACATGTCAAGTGCTTGTGTGATTTTCATAATGGACCAT
           ***********************************************************.

rb_4659    ATGCGAAAGAAGTCGGTGGAGCAGAAGCTGGCTACCTCAGGGGAGGGTCATGAATGGGGT
rb_49361   ATGCGAAAGAAGTCGGTGGAGCAGAAGCTGGCTACCTCAGGGGAGGGTCATGAATGGGGT
rb_103035  ATGCGAAAGAAGTCGGTGGAGCAGAAGCTGGCTACCTCAGGGGAGGGTCATGAATGGGGT
           ************************************************************

rb_4659    TTGCTGTTGGGCTTTGGTCCTGGTCTCACTTGTGAGACGGTCGTATTGAGAAGTGTGCCC
rb_49361   TTGCTGTTGGGCTTTGGTCCTGGTCTCACTTGTGAGACGGTCGTATTGAGAAGTGTGCCC
rb_103035  TTGCTGTTGGGCTTTGGTCCTGGTCTCACTTGTGAGACGGTCGTATTGAGAAGTGTGCCC
           ************************************************************

rb_4659    CTTGCCTCCCAATAG
rb_49361   CTTGCCTCCCAATAG
rb_103035  CTTGCCTCCCAATAG
           ***************
```
