## Additional File 4 for "Organ-Specific Long-Read Transcriptome Assembly of *Stenochlaena palustris* and Annotation of Anthocyanin Biosynthesis Genes"

```
CLUSTAL format alignment by MAFFT (v7.511)


rb_7618     ATGCCGGTCCCTGCTGGTGTGACTCTCCCCCCTTGCGCGCGCAAGATGGAGCGTGCTGAG
rb_74283    ATGCCGGTCCCTGCTGGTGCTACTTTCCCCCCTTGCGCGCGCAAGATGGAGCGTGCTGAG
rb_15005    ATGCCGGTCCCTGCTGGTGCTACTTTCCCCCCTTGCGCGCGCAAGATGGAGCGTGCTGAG
rb_52876    ATGCCGGTCCCTGCTGGTGCTACTTTCCCCCCTTGCGCGCGCAAGATGGAGCGTGCTGAG
            *******************. ***.***********************************

rb_7618     GGGCCTGCAACGGTTTTGGCGATCGGAACGGCGAATCCGCCCAATGTGTTCGAGCAGAGC
rb_74283    GGGCCTGCAACGGTTTTGGCGATCGGAACGGCGAATCCGCCCAATGTGTTCGAGCAGAGC
rb_15005    GGGCCTGCAACGGTTTTGGCGATCGGAACGGCGAATCCGCCCAATGTGTTCGAGCAGAGC
rb_52876    GGGCCTGCAACGGTTTTGGCGATCGGAACGGCGAATCCGCCCAATGTGTTCGAGCAGAGC
            ************************************************************

rb_7618     ACTTACCCTGATTTCTACTTCAACATCACCAACAGCAACCACATGACTGATCTCAAGGAG
rb_74283    ACTTACCCTGATTTCTACTTCAACATCACGAACAGCAACCACATGACTGATCTCAAGGAG
rb_15005    ACTTACCCTGATTTCTACTTCAACATCACCAACAGCAACCACATGACTGATCTCAAGGAG
rb_52876    ACTTACCCTGATTTCTACTTCAACATCACCAACAGCAACCACATGACTGATCTCAAGGAG
            ***************************** ******************************

rb_7618     AAGTTCCAGCGCATGTGTGATAAATCTGGGATTAAGAAGAGGTATATGTACCTGAATGAG
rb_74283    AAGTTCCAGCGCATGTGTGATAAATCTGGGATTAAGAAGAGGTATATGTACCTGAATGAG
rb_15005    AAGTTCCAGCGCATGTGTGATAAATCTGGGATTAAGAAGAGGTATATGTACCTGAATGAG
rb_52876    AAGTTCCAGCGCATGTGTGATAAATCTGGGATTAAGAAGAGGTATATGTACCTGAATGAG
            ************************************************************

rb_7618     GAAATCTTGAAGGCCAACCCAAGCATGTGTGCATACTGGGAGAAATCTTTAGATGTGAGG
rb_74283    GAAATCTTGAAGGCCAACCCAAGCATGTGTGCATACTGGGAGAAATCTTTAGATGTGAGG
rb_15005    GAAATCTTGAAGGCCAACCCAAGCATGTGTGCATACTGGGAGAAATCTTTAGATGTGAGG
rb_52876    GAAATCTTGAAGGCCAACCCAAGCATGTGTGCATACTGGGAGAAATCTTTAGATGTGAGG
            ************************************************************

rb_7618     CAAGATATGGTGGTTGTGGAGGTGCCAAAGCTCGGAAAGGAAGCGGCAGCCAAGGCGATC
rb_74283    CAAGATATGGTGGTTGTGGAGGTGCCAAAGCTCGGAAAGGAAGCGGCAGCCAAGGCGATC
rb_15005    CAAGATATGGTGGTTGTGGAGGTGCCAAAGCTCGGAAAGGAAGCGGCAGCCAAGGCGATC
rb_52876    CAAGATATGGTGGTTGTGGAGGTGCCAAAGCTCGGAAAGGAAGCGGCAGCCAAGGCGATC
            ************************************************************

rb_7618     AAGGAGTGGGGGCAGCCAAAGTCCAAGATCACGCACCTCATCTTCTGCACCACGAGTGGT
rb_74283    AAGGAGTGGGGGCAGCCAAAGTCCAAGATCACGCACCTCATCTTCTGCACCACGAGTGGT
rb_15005    AAGGAGTGGGGGCAGCCAAAGTCCAAGATCACGCACCTCATCTTCTGCACCACGAGTGGT
rb_52876    AAGGAGTGGGGGCAGCCAAAGTCCAAGATCACGCACCTCATCTTCTGCACCACGAGTGGT
            ************************************************************

rb_7618     GTAGACATGCCGGGTGCGGATTGGGCAATTACGAAGCTGCTGGGGCTGCGCCCCAGTGTG
rb_74283    GTAGACATGCCGGGTGCGGATTGGGCAATTACGAAGCTGCTGGGGCTGCGCCCCAGTGTG
rb_15005    GTAGACATGCCGGGTGCGGATTGGGCAATTACGAAGCTGCTGGGGCTGCGCCCCAGTGTG
rb_52876    GTAGACATGCCGGGTGCGGATTGGGCAATTACGAAGCTGCTGGGGCTGCGCCCCAGTGTG
            ************************************************************

rb_7618     AAGAGGCTGATGATGTACCAGCAGGGGTGCTTTGCTGGGGGCACAGTCATGAGGGTTGCA
rb_74283    AAGAGGCTGATGATGTACCAGCAGGGGTGCTTTGCTGGGGGCACAGTCATGAGGGTTGCA
rb_15005    A-----------------------------------------------------------
rb_52876    AAGAGGCTGATGATGTACCAGCAGGGGTGCTTTGCTGGGGGCACAGTCATGAGGGTTGCA
            *

rb_7618     AAAGACTTGGCTGAAAACAACAAGGGAGCCAGAGTATTGGTTGTGTGCAGTGAACTAACC
rb_74283    AAAGACTTGGCTGAAAACAACAAGGGAGCCAGAGTATTGGTTGTGTGCAGTGAACTAACC
rb_15005    ------------------------------------------------------------
rb_52876    AAAGACTTGGCTGAAAACAACAAGGGAGCCAGAGTATTGGTTGTGTGCAGTGAACTAACT


rb_7618     GCTGTGACATTCCGAGGGCCGAGCGACACACATCTAGACAGTCTTGTTGGGCAGGCTCTG
rb_74283    GCTGTGACATTCCGAGGGCCGAGCGACACGCATCTAGACAGTCTTGTTGGACAGGCTCTG
rb_15005    ------------------------------------------------------------
rb_52876    GCTGTGACATTCCGAGGGCCGAGCGACACGCATCTA------------------------


rb_7618     TTTGGGGACGGTGCCTCAGCTATCATCGTAGGCGCCGATCCCATCCCAGAAGTAGAGAGA
rb_74283    TTTGGGGACGGTGCCTCAGCTATCATCGTAGGCGCCGATCCCATCCCAGAAGTAGAGAGA
rb_15005    ------------------------------------------------------------
rb_52876    ------------------------------------------------------------


rb_7618     CCCTGGTTTGAGATTCACTATGTGGCATCCAACATCTTGCCGGACAGTGATGGCGCCATT
rb_74283    CCCTGGTTTGAGATTCACTATGTGGCATCCAACATCTTGCCGGACAGTGATGGCGCCATT
rb_15005    ------------------------------------------------------------
rb_52876    ------------------------------------------GAGAGTGATGGCGCCATT


rb_7618     GATGGACATCTCCGCGAGGTCGGCCTCACATTCCATCTCATGAAGGATGCACCCGAGCGA
rb_74283    GATGGACATCTCCGCGAGGTCGGCCTCACATTCCATCTCATGAAGGATGTACCCGGG-AT
rb_15005    ------------------------------------------------------------
rb_52876    GATGGACATCTCCGCGAGGTCGGCCTCACATTCCATCTCATGAAGGATGTACCTGGG-AT


rb_7618     CATCTCCAGAGACATAGGCATGGTGAGCATGCCTTTGAGGCCTTGGGATGA---------
rb_74283    CATCTCCAAGAACATAGGCACTGTGCTAAAGGATGCCTTTGAGAAGGTCTTTGGCAATGA
rb_15005    -----------------------------------------AGAAGGTCTTTGGCAATGA
rb_52876    CATCTCCAAGAACATTGGCACTGTGCTAAAGGATGCCTTTGAGAAGGTCTTTGGCAATGA


rb_7618     ------------------------------------------------------------
rb_74283    GAATGGAGAGCTCCCGAGCTACAATGATGTGTTCTGGATTGCCCACCCTGGTGGTCCTGC
rb_15005    GAATGGAGAGGTTCCGAGCTACAATGATGTGTTCTGGATTGCCCACCCTGGTGGTCCTGC
rb_52876    GAATGGAGAGGTTCCGAGCTACAATGATGTGTTCTGGATTGCCCACCCTGGTGGTCCTGC


rb_7618     ------------------------------------------------------------
rb_74283    TATTCTGGATCAAGTAGAACAGAAGCTGCAGCTGA-------------------------
rb_15005    TATTCTGGATCAAGTAGAGCAGAAGCTGCAGCTGAAACCAGAGAAGATGGCTCCGAGCCG
rb_52876    TATTCTGGATCAAGTAGAGCAGAAGCTGCAGCTGAAACCAGAGAAGATGGCTCCGAGCCG


rb_7618     ------------------------------------------------------------
rb_74283    ------------------------------------------------------------
rb_15005    ACAGGTATTGTCCGACTACGGCAACATGTCGAGTGCCTGTGTGCTCTTCATCATGGACCA
rb_52876    ACAGGTATTGTCCGACTACGGCAACATGTCGAGTGCCTGTGTGCTCTTCATCATGGACCA


rb_7618     ------------------------------------------------------------
rb_74283    --------------------ACCAGAATCTAGCCACCTCAGGAGAGGGCCATGAGTGGGG
rb_15005    TATGCGGAAGAAGTCGGTGGAGCAGAAGCTAGCCACCTCAGGAGAGGGCCATGAGTGGGG
rb_52876    TATGCGGAAGAAGTCGGTGGAGCAGAAGCTAGCCACCTCAGGAGAGGGCCATGAGTGGGG


rb_7618     ------------------------------------------------------------
rb_74283    ATTGCTCCTGGGTTTTGGCCCGGGTCTTACCTGTGAGACAGTTGTATTAAGAAGTGTGCC
rb_15005    ATTGCTCCTGGGTTTTGGCCCGGGTCTTACCTGTGAGACAGTTGTATTAAGAAGTGTGCC
rb_52876    ATTGCTCCTGGGTTTTGGCCCGGGTCTTACCTGTGAGACAGTTGTATTAAGAAGTGTGCC


rb_7618     ----------------------
rb_74283    CCTTGCTACTGCTACTGCATAG
rb_15005    CCTTGCTACTGCTACTGCATAG
rb_52876    CCTTGCTACTGCTACTGCATAG
```
