## Additional File 5 for "Organ-Specific Long-Read Transcriptome Assembly of *Stenochlaena palustris* and Annotation of Anthocyanin Biosynthesis Genes"

CLUSTAL format alignment by MAFFT (v7.511)

```
MK920230.1 ------------------------------------------------------------
rb_103797  ATGTTGTGGCTCTTTGTTGCCATAATAAGAATAAGAGAACATGGAGAGGAGGAGCATGAG
rb_54792   ------------------------------------------------------------


MK920230.1 ---------------------------------------------------------ATG
rb_103797  TTTGCGCAGAGCACATTTTACGAAGCAGAGGCGGAGCTGCCAGCCATTGCCATCCTAATG
rb_54792   ---------------------------------------------------------ATG
                                                                    ***

MK920230.1 G---ACAAGCCCCTTCACTCCACAGTCTTGGTTACAGGCTGCACCGGCCACATCGGCTCT
rb_103797  GCCCCCAAATCCCCTCATTCTACCGTCTTGGTCACTGGCTCCACGGGCCACCTCGGCTCT
rb_54792   GCCCCCAAATCCCCTCATTCTACCGTCTTGGTCACTGGCTCCACGGGCCACCTCGGCTCT
           *    ***..***.***.**.** ********.** **** *** ****** ********

MK920230.1 TGGCTCGTCTTACGCCTCCTCGAGAAGGGCTACAGCGTCAGAGCTGCCGTCCTTGATCCG
rb_103797  TGGCTCGTCCTCCGCCTCCTCGAGAAGGGCTACTTCGTCAGAGCTGCTGTCCTTGACCCT
rb_54792   TGGCTCGTCCTCCGCCTCCTCGAGAAGGGCTACTTCGTCAGAGCCGCTGTCCTTGACCCT
           *********.* *********************  *********.**.********.**

MK920230.1 GAAGACAAGCAAGATGTGCAACCCCTCCTTGGCCTCGCTCCATCGTTGGCCGAGAGACTT
rb_103797  GAAGACAAGCAAGATCTGCAACCCCTCCTTAGCTTCGCTCCGGCTTCAACTGAGAGACTT
rb_54792   GAAGACAAGCAAGATCTGCAACCCCTCCTTAGCTTTGCTCCGGCTTCAACTGAGAGACTT
           *************** **************.**.*.*****. * *...*.*********

MK920230.1 GAAATTGTGAAAGCAGACCTCCGTGTCAAAGGGGACTTCGACAAAGTTGCCCAAGGGTGT
rb_103797  GAAATTATGAAAGCAGACCTCCGTGTGGAAGGCGACTTTGACAAAGTTGTGGAAGGGTGT
rb_54792   GAAATTATGAAAGCAGACCTCCGTGTGGAAGGCGACTTTGACAAAGTTGTGGAAGGGTGT
           ******.******************* .**** *****.**********.  ********

MK920230.1 GATGGTGTTTTTCACTTGGCATGTCCAACCAATTTTCTTGTAGAAGATCCCCAGAAGGAT
rb_103797  GAGGGGGTTTTTCACTTGGCATGTCCGACCGGTTTTATGGTAGAGGATCCCCAGAAGGAT
rb_54792   GAGGGGGTTTTTCACTTGGCATGTCCGACCGGTTTTATGGTAGAGGATCCCCAGAAGGAT
           ** ** ********************.***..**** * *****.***************

MK920230.1 GTGATTGAACCGGCAGTTCAAGGCACTTTGAATGTGCTGAATGCTTGTGTCAAGTCTTCT
rb_103797  GTGATTGAACCGGCAGTTCAAGGCACCCTAAATGTGCTGAATGCTTGTGTGAAATCTTCT
rb_54792   GTGATTGAACCGGCAGTTCAAGGCACCCTAAATGTGCTGAATGCTTGTGTGAAATCTTCT
           **************************..*.******************** **.******

MK920230.1 TCAGTTAAGCGTGTGATACATGTGTCATCTGCGGCCGCTATCCGTTTCTCAGGGAAGGAG
rb_103797  TCTGTGAAGCGTGTGATACATGTGTCATCTGCGGCTGCTATCCGGTTCTCAGGCAAGGAG
rb_54792   TCTGTGAAGCGTGTGATACATGTGTCATCTGCGGCTGCTATCCGGTTCTCAGGCAAGGAG
           ** ** *****************************.******** ******** ******

MK920230.1 GTGGCAGAGCAAGTGTTTGATGAGTCATGCTGGACAGATGTGGACTTTTGTATCGACAAT
rb_103797  GTGGCAGAGCAAGTGTTTGATGAGTCATGCTGGACGGACGTGGACTTCTGCATCAAAAAT
rb_54792   GTGGCAGAGCAAGTGTTTGATGAGTCATGCTGGACGGACGTGGACTTCTGCATCAAAAAT
           ***********************************.**.********.**.***.* ***

MK920230.1 AAAATACCTGGATGGCCATACTTTGTGGGCAAAACGCTAGGGGAGAAAGCTGCGGTGGAG
rb_103797  AAAATACCTGGATGGCCTTACTTTGTGGGCAAGACATTAGGGGAGAAAGCGGCGGTGGAG
rb_54792   AAAATACCTGGATGGCCTTACTTTGTGGGCAAGACATTAGGGGAGAAAGCGGCGGTGGAG
           ***************** **************.**..************* *********

MK920230.1 TTTGCCAAAGAACACAACCTGGACTTGGTAGTTGTCAATCCAAGTATTGTACATGGACCT
rb_103797  TTTGCAGAAAAGCACAACATGGACTTGGTAGTTGTCAATCCGGGCATTGTAAATGGACCA
rb_54792   TTTGCAGAAAAGCACAACATGGACTTGGTAGTTGTCAATCCGGGCATTGTAAATGGACCA
           ***** .**.*.****** **********************..*.****** *******

MK920230.1 TTCTTGCTTTCCAACATTCCCAACAGCGTAAAGGATTGCCTGGCTCTAATAACAGGGGAG
rb_103797  TTCTTGCTTTCCAAGATTCCCAATAGCATAAAGGATTGCCTGGCTTTAATAACAGGGGAG
rb_54792   TTCTTGCTTTCCAAGATTCCCAATAGCATAAAGGATTGCCT-------------------
           ************** ********.***.*************

MK920230.1 CTGTTCAAGCTGCCCTTTTTAGCGCGCATGTCATACATCCACACAGATGACTTGGTGCGA
rb_103797  CTGTTCAAGCTGCCTTTTACGCAACGCATGTCGTACGTTCACACAGATGACCTGGTGCGA
rb_54792   ------------------------------------------------------------


MK920230.1 GCCTTCATTTTTCTTTATGAAAATCCTGAGGCGAGTGGCCGCTATATCTGTTCTGCAGTT
rb_103797  GCCTTCATTTTTCTTTATGAAAATCCTGAGGCGAGTGGCCGCTATATCTGTTCAGCAGTT
rb_54792   ------------------------------------------------------------


MK920230.1 GATGCCTCCATCACAGAGGTGGTCAATATCATCTCTTCTCGCTATCCCGAATTCAAAATT
rb_103797  GATGCCTCCATCACAGAAGTTGTCAAAATCATCTCTTCTCACTATCCAGAATTTAAAATT
rb_54792   -------CCATCACAGAAGTTGTCAAAATCATCTCTTCTCACTATCCAGAATTTAA----
                  **********.**.***** *************.****** *****.**

MK920230.1 CCCACAGATCTGGCCTACCTTGGGGAGCCTGTTATTCATTATTTCGACTCTTCCAAGTTG
rb_103797  CCCACAGACCTGGCCTACCTTGGGGAGCCCTTCATTCATTATTTTGACTCTTCCAAGTTG
rb_54792   ------------------------------------------------------------


MK920230.1 AAGAGGCTTGGCTTTCAGTTTAACTTCACACTCGAAGACATGTACCATGATGCCATCAAA
rb_103797  AAGAGGCTTGGCTTCCAGTATAATTTCACACTCGAAGATATGTACCGTGATGCCATTGAA
rb_54792   ------------------------------------------------------------


MK920230.1 TGCTGCAAGGAGAAAGGGCTGTTATAA---------------------------------
rb_103797  GGCTGCAAGGAAAAAGGACTCTTACAAGCCTTTGAAAGCGCTCCTCTTCAGGCGCACTAG
rb_54792   ------------------------------------------------------------
```
