## Additional File 6 for "Organ-Specific Long-Read Transcriptome Assembly of *Stenochlaena palustris* and Annotation of Anthocyanin Biosynthesis Genes"

```
CLUSTAL format alignment by MAFFT (v7.511)


rb_13889   ------------------------------------------------------------
MK920231.1 ATGGCTCCTAATGCTGTCGCAGTTATGGCCTTGCCAGAGTTAGTGTGTGTAACGGGAGCC


rb_13889   ------------------------------------------------------------
MK920231.1 TCAGGCCATCTGGGTACCTGGCTTGTGAAGCGCTTGCTTGAGAGAGGGTATAGAGTGCGT


rb_13889   ------------------------------------------------------------
MK920231.1 GCCACGGTCAGAAACCCAGGGGACCCAACGAAAACGGCCATTTTAAGAAATCTTCCTGGA


rb_13889   ------------------------------------------------------------
MK920231.1 GCTGATGAGAGGCTGCAAATTGTTAGAGGCCAGCTTCTTGAAGAGAGTGGGTTTGATGAT


rb_13889   ------------------------------------------------------------
MK920231.1 GCCGTGAATGGCTGTGATGGAGTTTTCCATTGTGCTTGCCCCACTGAATTTTCCTTTGAG


rb_13889   ------------------------------------------------------------
MK920231.1 GACCCCTATAAGGACTGCATTGAGCCAGCTGTGAATGGGACTGTGAATGTGCTAAAGGCC


rb_13889   ------------------------------------------------------------
MK920231.1 TGCGCCAAGGCCCCCTCTATCAAGCGGATCGTCTTCACATCATCTGCAGCAGCAGTGAGG


rb_13889   ------------------------------------------------------------
MK920231.1 TTTACGCAGCTGGAGGATATTGCGGGACAGCGATTCGATGAGAACTGTTGGACAGACATT


rb_13889   -------------------------ATGGATGGCCCTACTTTGTATCCAAGGCACTCTCT
MK920231.1 GAGATGTGCGAGAGAGACAAGCCTCATGGATGGCCTTACTTCGTTTCCAAGACCCTCTCT
                                    **********.*****.** ******.* ******

rb_13889   GAGAAGAAAGCATTGGAGCTTGCACCAAGAATACAAGCTAGATTTGGTCACAATATTGCC
MK920231.1 GAAAAGAAAGCATTTGAGCTTGCA-CAAGAATACAACCTGGATTTGGTCACAATATTGCC
           **.*********** ********* *********** **.********************

rb_13889   AGTGTTAGTGAATGGCCCCTTTCTAATTAACAAAATTCCCAATAGTGTAGCAGATGCGAT
MK920231.1 ACCCTTGGTCAATGGCCCCTTTCTAATTGATAAAATTCCCAACAGTGTAGCAGATGCAAT
           * . **.** ******************.*.***********.**************.**

rb_13889   CTCCCTTGTCACAGGTGATACTTCCCATCACAAATTTATTCGATGGATCTCATATGTACA
MK920231.1 CTCCCTTGTCACAGGCGATATCTCTCATCACAAATTCATTCGTCGGATCTCATATGTACA
           ***************.****..**.***********.***** .****************

rb_13889   CATTGATGACATCATTGATGCACACATCTTCTTATATGAGACACCTACCGCTCAAGGTCG
MK920231.1 CATTGATGACATCATTGATGCACACATCTTCTTGTACGAGACACCTGCTGCTCAAGGTCG
           *********************************.**.*********.*.***********

rb_13889   CTATATTGGGTCCGCTGTGGATGCATCCATCATGGAAGTTGCTGAGCTCCTCAAGAAGCT
MK920231.1 TTACAATGGGTCCTGTGCAGATGCATCCATCACAGAGATTGCTGAACTCCTCAAGAAGCT
           .**.* *******  **..*************..**..*******.**************

rb_13889   CTTCCCCAAGATCAAGCTACCGGATAATTTTGACTACGTGGGGGAGATTGTTCCTCACTA
MK920231.1 CTTCCCCAAGATCAAGCTACCGGATAACTTTGATTATGTGGGAGAGATACTGCTCCATTA
           ***************************.*****.**.*****.*****  * *..**.**

rb_13889   CATGGATAATTCAAAGCTAAAAGATCTGGGCTTTGAGTTCAAACATGGGATGGTTGACAT
MK920231.1 CATGGACAATTCGAAGCTGAAAAAGTTAGGCTTCGAGTTCAAACACGGGCTGGCCGACAT
           ******.*****.*****.***.* .*.*****.***********.*** ***..*****

rb_13889   GTTTGAAGGCGCAGTCCAATGTTGTGTCGAGAAGGGGCTCCTCCAACATCCAGAACAA--
MK920231.1 GTTTGAGAGTGCAGTCAAATGCTGTGTGGAAAAAGGTCTGCTCCAATATCCTGGAGAAGC
           ******..*.****** ****.***** **.**.** ** ******.**** *.* **

rb_13889   ----TAA
MK920231.1 CCCTTGA
               *.*
```
