## Additional File 7 for "Organ-Specific Long-Read Transcriptome Assembly of *Stenochlaena palustris* and Annotation of Anthocyanin Biosynthesis Genes"

```
CLUSTAL format alignment by MAFFT (v7.511)

rb_10724   ------------------------------------------------------------
rb_11399   ATGGAGGCTTGCCTTAATGAGAAGGGAATTAGGGAGAGGGGAAGAGTGTGTGTGACTGGA
rb_57372   ------------------------------------------------------------
MK920232.1 ATGGAG-------------------------GGTGGAAATGGCGAGTGTGTGTAACGGGG


rb_10724   ------------------------------------------------------------
rb_11399   GCATCCGGCTACCTGGCATCCTGGCTTGTCATGCGCTTGCTCCAACGAGGCTACACCGTG
rb_57372   ------------------------------ATGCGCTTGCTCCAACGAGGCTACACCGTG
MK920232.1 GCATCAGGCTACCTGGGATCCTGGCTTGTCATGCGCTTGCTCCAACGAGGCTACTCTGTG


rb_10724   ------------------------------------------------------------
rb_11399   CGAGGCACCGTTCGCAACTTAGAAGACAAGTCAAAGACCGATCACCTTCTCTCACTACCA
rb_57372   CGAGGCACCGTTCGCAACTTAGAAGACAAGTCAAAGACCGATCACCTTCTCTCACTACCA
MK920232.1 CGAGGCACCGTTCGCAGCCTAGAAGACAAGTCAAAGACAGAGCATCTTCTCTCACTACCC


rb_10724   ------------------------------------------------------------
rb_11399   GAGGCAAAGGAGAAGCTAGAGTTAGTGGAAGCAGATATCTTAGATGAGTCTTCACTGCAT
rb_57372   GAGGCAAAGGAGAAGCTAGAGTTAGTGGAAGCAGATATCTTAGATGAGTCTTCACTGCAT
MK920232.1 GAGGCAAAGGAGAAGCTAGAGCTGGTGGAAGCAGATTTGTTAGTTTATTCTTCAATTCCC


rb_10724   ------------------------------------------------------------
rb_11399   GTTGCAGTCAAAGGTTGTGTTGGGGTATTCGCGATTGCCGCACCTTTAATTGACCCCTCA
rb_57372   GTTGCAGTCAAAGGTTGTGTTGGGGTATTCGCGATTGCCGCACCTTTAATTGACCCCTCA
MK920232.1 GCCGCAGTGAAGGGTTGTGATGGGGTATTCGCTGTTGCTGTACCTATTCTTGACCCCTCA


rb_10724   ---------------ATGATAGATTTGACCATGGCATCCACTATGAACCTATTGAGAGCA
rb_11399   AATGACCAGAGTTCAATGATAGATTTGACCATGGCATCCACTATGAACCTATTGAGAGCA
rb_57372   AATGAC------------------------------------------------------
MK920232.1 AAAGACCAGAGCGGGATGATAGAGTTGGCCATGACATCCACAATGAACCTACTAAGAGCA


rb_10724   TGTCATGAGTCACATTGTGTAAAGAGGGTGGTCATGACTTCATCTTACTATGCCGCTATT
rb_11399   TGTCATGAGTCACATTGTGTAAAGAGGGTGGTCATGACTTCATCTTACTATGCCGCTATT
rb_57372   ------------------------------------------------------------
MK920232.1 TGCGATGAGTCACATTGCGTAAAGAGGGTGGTCATGACTTCAACTTACTACAGTGCTATT


rb_10724   CAGGTCCCTCACGAAGGGGAGATTCCGTTTGAAGTGGACGAGTCCTCATGGAGTTCTGTG
rb_11399   CAGGTCCCTCACGAAGGGGAGATTCCGTTTGAAGTGGACGAGTCCTCATGGAGTTCTGTG
rb_57372   CAGGTCCCTCACGAAGGGGAGATTCCGTTTGAAGTGGACGAGTCCTCATGGAGTTCTGTG
MK920232.1 CAGGTTCCACATGAAGGGGAGATTCCATTTGAAGTAGATGAATCCTCGTGGACCTCCGTG
           *****.** **.**************.********.**.**.*****.**** .**.***

rb_10724   GAGTACTTAAGGCACAAGAAACTATCATGTTGGGGCTACATGGTTGCCAAAACCATGGCA
rb_11399   GAGTACTTAAGGCACAAGAAACTATCATGTTGGGGCTACATGGTTGCCAAAACCATGGCA
rb_57372   GAGTACTTAAGGCACAAGAAACTATCATGTTGGGGCTACATGGTTGCCAAAACCATGGCA
MK920232.1 GAGTATTGTATGCAGCAAAAGATGTCATGTTGGGGTTACATGGTTGCAAAAACCTTGGCA
           *****.*  * ***  *.**. *.***********.*********** ****** *****

rb_10724   GAGAAGGCAGCATTTGAGTTTGCAGAAGAGAATGGCTTGGATTTTATTTCCATTGCGCCA
rb_11399   GAGAAGGCAGCATTTGAGTTTGCAGAAGAGAATGGCTTGGATTTTATTTCCATTGCGCCA
rb_57372   GAGAAGGCAGCATTTGAGTTTGCAGAAGAGAATGGCTTGGATTTTATTTCCATTGCGCCA
MK920232.1 GAGAAAGCAGCATTTGAGTATGCAAAAGAGAAAGAGTTCGATCTGATTTCCATCTTACCA
           *****.************* ****.******* *. ** ***.* ********. ..***

rb_10724   TCACTTGTTGGGGGGCCCTTTCTCACACCTTCTATTCCATCAAGTGTCTCGCTATTTCTA
rb_11399   TCACTTGTTGGGGGGCCCTTTCTCACACCTTCTATTCCATCAAGTGTCTCGCTATTTCTA
rb_57372   TCACTTGTTGGGGGGCCCTTTCTCACACCTTCTATTCCATCAAGTGTCTCGCTATTTCTA
MK920232.1 TCACTTGTGGGGGGGGCCTTTTTAACGCCTTCTGTTCCAGGGAGCGTCGCACTTTACTTG
           ******** ****** *****.* **.******.*****  .**.*** *.** * ..*.

rb_10724   GCCACCATGATTGGTACCACAGAGTGGATGATCCCTTTAATGGGATACATGTCATATGTG
rb_11399   GCCACCATGATTGGTACCACAGAGTGGATGATCCCTTTAATGGGATACATGTCATATGTG
rb_57372   GCCACCATGATTGGTACCACAGAGTGGATGATCCCTTTAATGGGATACATGTCATATGTG
MK920232.1 GGCGCCATGACTGATGTGGCGACGTGGATGAGCTCTCTACTTGGTTACATGTCGTATGTG
           * *.******.**.*.. .*.. ******** *.**.** * ** ********.******

rb_10724   CATGTGGATGATGTTATTCAAGCTCATATCTTTTTGATGGAGGAATCACATGCAAAAGGG
rb_11399   CATGTGGATGATGTTATTCAAGCTCATATCTTTTTGATGGAGGAATCACATGCAAAAGGG
rb_57372   CATGTGGATGATGTTATTCAAGCTCATATCTTTTTGATGGAGGAATCACATGCAAAAGGG
MK920232.1 CATGTGGATGATACTATACAAGCTCATATATTATTGATGGAGGACTCACATGCAAAAGGG
           ************..*** *********** ** *********** ***************

rb_10724   CGGTACCTTTGTTCTGCTGTGGATGGTTCCCTTAGTGATGTGCTTCATGAGGTCTTACCT
rb_11399   CGGTACCTTTGTTCTGCTGTGGATGGTTCCCTTAGTGATGTGCTTCATGAGGTCTTACCT
rb_57372   CGGTACCTTTGTTCTGCTGTGGATGGTTCCCTTAGTGATGTGCTTCATGAGGATCGTCTC
MK920232.1 CGCTATCTTTGCTCCGCCGTGGATGCAACATTGGATGATGTGCTTCATGAGGTGTTACCT
           ** **.*****.**.**.*******   * .* ..*****************

rb_10724   CTTCTTGAGTCTTTTAATATTCCTCCACCAAAAGCGATCGTCTCTCAAGACAAGGATAAA
rb_11399   CTTCTTGAGTCTTTTAATATTCCTCCACCAAAAGCGATCGTCTCTCAAGACAAGGATAAA
rb_57372   CTCAAGACAAGGATAA--------------------------------------------
MK920232.1 CTTCTCAAGTCTTTTAATATCCCTCCACCAAAAGTGGCTGCCTCTCTCGATGAAGACCCA


rb_10724   TCAAAGGCCTACTACAAACTCAACACAAATAAACTCAAAAACTTGGGCTTCACTTATAAA
rb_11399   TCAAAGGCCTACTACAAACTCAACACAAATAAACTCAAAAACTTGGGCTTCACTTATAAA
rb_57372   ------------------------------------------------------------
MK920232.1 ACAAAAGCTTACTACAAAATCAACACAACAAAACTCAAAGACATGGGATTCACTTATGAA


rb_10724   TTTGACACATTGCAAAAAATTTTCAAAGATGCTATTCTCTTCTTAATTGAGAAAGGTTAC
rb_11399   TTTGACACATTGCAAAAAATTTTCAAAGATGCTATTCTCTTCTTAATTGAGAAAGGTTAC
rb_57372   ------------------------------------------------------------
MK920232.1 TTTGACACACTCCAGAAAATTTTTAAAGATTCAGTGCTTTTCTTCCTTGAAAAAGGCCAC


rb_10724   TTGAAGCAACAACAAAAAGGTGTTGCCTTGTAA
rb_11399   TTGAAGCAACAACAAAAAGGTGTTGCCTTGTAA
rb_57372   ---------------------------------
MK920232.1 TTAAAGCAAGTACAATAG---------------
```
